# TEMPO: A system to sequentially label and genetically manipulate vertebrate cell lineages

**DOI:** 10.1101/2021.10.27.466134

**Authors:** Isabel Espinosa-Medina, Daniel Feliciano, Carla Belmonte-Mateos, Jorge Garcia-Marques, Benjamin Foster, Rosa Linda Miyares, Cristina Pujades, Minoru Koyama, Tzumin Lee

**Affiliations:** Janelia Research Campus, Howard Hughes Medical Institute; Ashburn, VA, 20147, USA; Department of Experimental and Health Sciences, Universitat Pompeu Fabra, PRBB, Barcelona, 08003, Spain; Centro Nacional de Biotecnologia, Consejo Superior de Investigaciones Cientificas, Madrid, 28049, Spain

## Abstract

During development, regulatory factors appear in a precise order to determine cell fates over time. To investigate complex tissue development, one should not just label cell lineages but further visualize and manipulate cells with temporal control. Current strategies for tracing vertebrate cell lineages lack genetic access to sequentially produced cells. Here we present TEMPO (Temporal Encoding and Manipulation in a Predefined Order), an imaging-readable genetic tool allowing differential labelling and manipulation of consecutive cell generations in vertebrates. TEMPO is based on CRISPR and powered by a cascade of gRNAs that drive orderly activation/inactivation of reporters/effectors. Using TEMPO to visualize zebrafish and mouse neurogenesis, we recapitulated birth-order-dependent neuronal fates. Temporally manipulating cell-cycle regulators in mouse cortex progenitors altered the proportion and distribution of neurons and glia, revealing the effects of temporal gene perturbation on serial cell fates. Thus, TEMPO enables sequential manipulation of molecular factors, crucial to study cell-type specification.

**One-Sentence Summary:** Gaining sequential genetic access to vertebrate cell lineages.

## Main Text

Cell specification during tissue and organ formation depends on spatial position, temporal patterning and cell-lineage relationships (*1-3*). In many biological systems the timing of proliferative events and the order of expression of differentiation promoting factors determine the cell’s anatomical distribution and identity. A well-studied example is the development of neuronal circuits in Drosophila, formed by multiple neuronal types assembled in a precise spatial and temporal manner. Neural progenitors express a cascade of transcription factors and differentiate following an invariant order, determining temporal patterns of functional circuit assembly (*4*). In more complex organisms, such as vertebrates, emergence of cell diversity during tissue formation is also regulated by spatial and temporal patterning. For example, gene expression cascades are essential for the progressive differentiation of pancreatic cell types, and for the sequential emergence of neuronal subtypes in the central nervous system (*5-6*). While the transcriptional timing of many differentiation genes has been established, the temporal boundaries in which these factors can promote cell fate determination have not been defined. This limits our ability to predict, and potentially correct, developmental defects caused by misregulation of cell fate regulatory genes. Thus, establishing the cellular origins and spatio-temporal interactions underlying complex tissue formation is crucial to determine the mechanisms of cell specification in development and disease and to eventually produce cell types at will for cell replacement therapy (*7*).

Genetic labeling and manipulation strategies which preserve the native tissue context are necessary to establish the link between cell origins, identity and spatial distribution. Existing technologies require live imaging to reconstruct cell histories at single-cell resolution (*8-9*). However, in most complex organisms, live imaging over long periods of time is limited by sample thickness and phototoxicity, impairing direct visualization of cell interactions and differentiation patterns. Thus, defining cell histories in complex organisms often requires retrospective analysis, that is inferring past developmental relationships or cell lineages from end point cell-specific labels (*10-12*). In organisms with stereotypic development such as Drosophila, complete cell histories can be inferred by combining partial reconstructions from multiple individuals (*13*). In contrast, in organisms without stereotypic development like vertebrates it is necessary to map entire cell lineages in a single sample and compare experiments to extract general principles due to the higher variability across individuals (*14*). Advances in synthetic biology and sequencing technologies have enabled the development of DNA recorders that allow retrospective and simultaneous analysis of both cell identity and lineages (*15*). However, these approaches lack spatial information and have low temporal resolution due to dissociation of tissues, cell loss and stochastic recorder depletion (*16-17*). In contrast, imaging-based strategies enable spatial identification of cell clones but lack the necessary number of labels to distinguish subsequent cell generations and infer their mitotic connections (*18*). We recently developed an imaging-based approach in Drosophila called CLADES, which allows differential labelling of sequentially produced neurons and simultaneous visualization of anatomical position and temporal lineage information in the same sample (*19*). However, this strategy relies on the genomic integration of multiple synthetic transgenes and controlled splicing designs which are not compatible with vertebrate transgenesis. Further, none of the current technologies allow genetic manipulation of vertebrate cell histories in a predefined order, limiting functional studies on the role of temporal cell-determination factors in vivo. Therefore, there is a need for a new genetic tool to simultaneously label and manipulate cell histories in vertebrates.

Here we present TEMPO, a new system based on imaging and CRISPR/Cas9 that allows temporal genetic labelling and manipulation of vertebrate cell histories in vivo, including zebrafish and mice. TEMPO relies on a compact transgene containing three different fluorescent reporters in mutually exclusive coding frames. A conditional cascade of gRNAs, along with Cas9, drive activation and inactivation of the reporters in a predefined order, labelling consecutive cell generations with different colors. By introducing cell-fate regulatory genes in different TEMPO coding frames, we generated defined temporal windows of genetic perturbations that allow the study of serial temporal fates in a single sample. Thus, TEMPO establishes a foundation to manipulate cell-determination factors in a specific temporal sequence while allowing simultaneous visualization of spatial and temporal cell relationships in vertebrate models.

## RESULTS

### Design and validation of TEMPO

An ideal system for temporal cell labelling and genetic manipulation in complex organisms should consist of: (1) a predefined and irreversible sequence of transgenes to distinguish and manipulate multiple cell generations, (2) a compact design to allow efficient genome integration and germline transmission and (3) a method compatible with available cell type targeting and imaging techniques to achieve spatio-temporal resolution.

To build such a versatile system in vertebrates, we repurposed the recently developed CaSSA switches (*20*). These genetic switches are disrupted reporter genes which become activated by a CRISPR/Cas9 double-strand break (DSB) between two direct repeats (homologous sequences), followed by single-strand annealing (SSA) repair mechanism that recombines both repeats resulting in a scarless sequence (Fig.1A) (*21*). In contrast to non-homologous repair mechanisms (*22*), SSA has predictable outcomes that are important for engineering conditional transgenes. To create a compact transgene expressing alternative reporter genes, we developed Frame switches (Fig.1B) by placing CaSSA reporters in different coding frames separated from each other by 2A peptide sequences. In this design, a preactivated reporter (ON) is placed downstream of an inactive CaSSA reporter (OFF). After Cas9 editing and SSA repair, the CaSSA reporter is activated resulting in a frame shift that inactivates the downstream reporter. To activate several of these reporters in a predefined order, we deployed a conditional guide RNA (gRNA) switch (*19*). Similar to CaSSA switches, a gRNA switch is a disrupted gRNA sequence that gets activated by another gRNA and Cas9 editing followed by SSA repair (Fig.1C). With these elements we engineered TEMPO, a compact tri-cistronic transgene containing three fluorescent protein sequences in three different frames sequentially activated by a parallel cascade of gRNAs and Cas9 nuclease. The reporter cascade starts with a preactivated CFP reporter in the first temporal window (T1), followed by the activation of RFP CaSSA reporter by gRNA-1 (T2), which also activates a gRNA-2 switch, to subsequently drive YFP CaSSA reporter activation (T3). Importantly, only one reporter is expressed at a time, each step in the cascade is irreversible and only repair through SSA allows reporter activation (Fig.1D). This design maximizes reporter diversity while reducing the number of transgenes needed to allow genetic access to spatio-temporal cell relationships (Fig.1E).

**Fig. 1.**
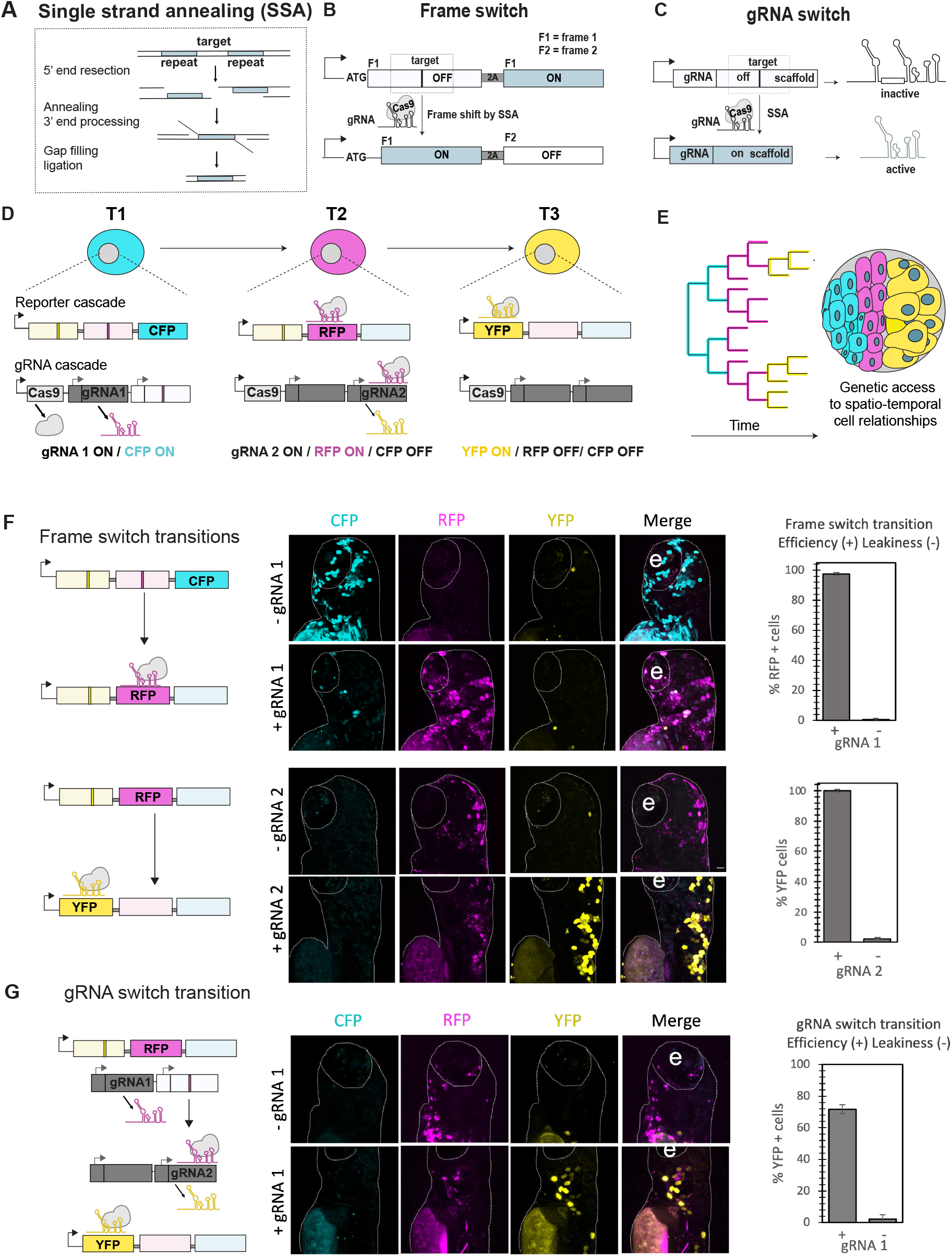
A conditional cascade of genetic switches to access temporal cell relationships in vivo. **(A)** Representation of the repair mechanism Single-Strand Annealing (SSA), required for activation of Frame and gRNA switches (B and C). **(B)** The frame switch design allows activation of a reporter (OFF---ON, F1=frame 1) after a double strand break (DSB) by CRISPR/Cas9 followed by SSA repair that causes a frame shift, inactivating the downstream reporter (OFF, F2=frame 2). **(C)** A conditional gRNA activated by another gRNA through SSA repair. **(D)** TEMPO transgenes enable the ordered activation of a tri-cistronic reporter sequence (CFP-RFP-YFP, T= temporal windows) by a conditional gRNA cascade. gRNA and their corresponding targets in the switch sequences are color coded. **(E)** TEMPO allows genetic access and simultaneous visualization of sequentially produced cells in their anatomical context. **(F)** Frame switch transitions assessed by expression of fluorescent reporters in 3 day-post-fertilization (dpf) zebrafish larvae. Plots indicate the mean SEM of the fluorescence for the reporter activated in the presence (+) or absence (-) of the corresponding gRNA, representing efficiency or leakiness of the transition, respectively (n=19 fish, 6 independent experiments). **(G)** A gRNA switch fluorescent assay enables characterization of the conditional gRNA-2 switch efficiency (+) and leakiness (-) in the presence or absence of gRNA-1, respectively (n=12 fish, 6 independent experiments). e= eye.

We independently characterized the activation of each Frame switch by injecting 1-cell-stage zebrafish embryos with integrative constructs ubiquitously expressing: (1) the TEMPO reporter with either CFP or preactivated RFP and (2) Cas9 and either gRNA-1 or gRNA-2 required for RFP or YFP activation, respectively. We found high efficiency of color transition (>90%) for both Frame switches in the presence of the corresponding gRNA, and minimal leakiness in its absence (Fig.1F). For the gRNA switch design, our previous version had ∼50% activation efficiency by SSA both in Drosophila and zebrafish (*19*). However, higher efficiency is desired to avoid limited progression of the cascade. To improve gRNA switch repair efficiency, we tested multiple variants with longer direct repeats in the scaffold region (Fig. S1). Based on previous works, we reasoned that SSA efficiency would increase with higher homology repeat lengths (*23*) and modifying this gRNA region should not affect gRNA structure or function (*24*). We tested these variants in zebrafish embryos and obtained a V2.0 gRNA switch with enhanced efficiency (>70%) and minimal leakiness (Fig. 1G). These results could also be generalized to other gRNA sequences (Fig.S2), demonstrating that rational design of conditional gRNA switches results in optimal performance and versatility of this system.

To validate the full TEMPO reporter cascade, we injected 1-cell-stage zebrafish embryos with two integrative constructs ubiquitously expressing: (1) the TEMPO reporter with the preactivated gRNA-1 and (2) Cas9 and the inactive gRNA-2 switch (Fig. 2A). The efficiency of co-expression of both plasmids was measured to be >88% (Fig. S3). We then monitored the progression of the reporter cascade by taking snapshots of the same embryos at 8 hours postfertilization (hpf), 24hpf, 48hpf, 72hpf and 96hpf. At 8hpf, most TEMPO+/Cas9+ cells expressed CFP and some started to transition to the second temporal reporter, RFP. By 24hpf most cells expressed RFP while the last cascade transition to YFP was apparent by 48hpf and became predominant by 72hpf (Fig.2B-E). Intermediate states of cells transitioning from one color to the next could be distinguished by the co-expression of two reporters (Fig. 2C). While most TEMPO+ cells were dividing epithelial cells that expressed the last YFP reporter at the end of the recording, we found that cells that did not divide in the same time period halted reporter transitions (Fig. 2C). This is consistent with SSA repair activity restricted to dividing cells (*21*). We defined the percentage of color transition (% CT) as the final proportion of TEMPO+ cells (Output) with respect to the proportion of initial TEMPO+ cells (Input). This revealed that >90% of total TEMPO+ cells transitioned to RFP and >85% reached the last color reporter, YFP, by the end of the recording (Fig. 2F). These results demonstrate that TEMPO works efficiently and the reporters are expressed in a predefined order, allowing spatio-temporal labelling of cell histories in live zebrafish.

**Fig. 2.**
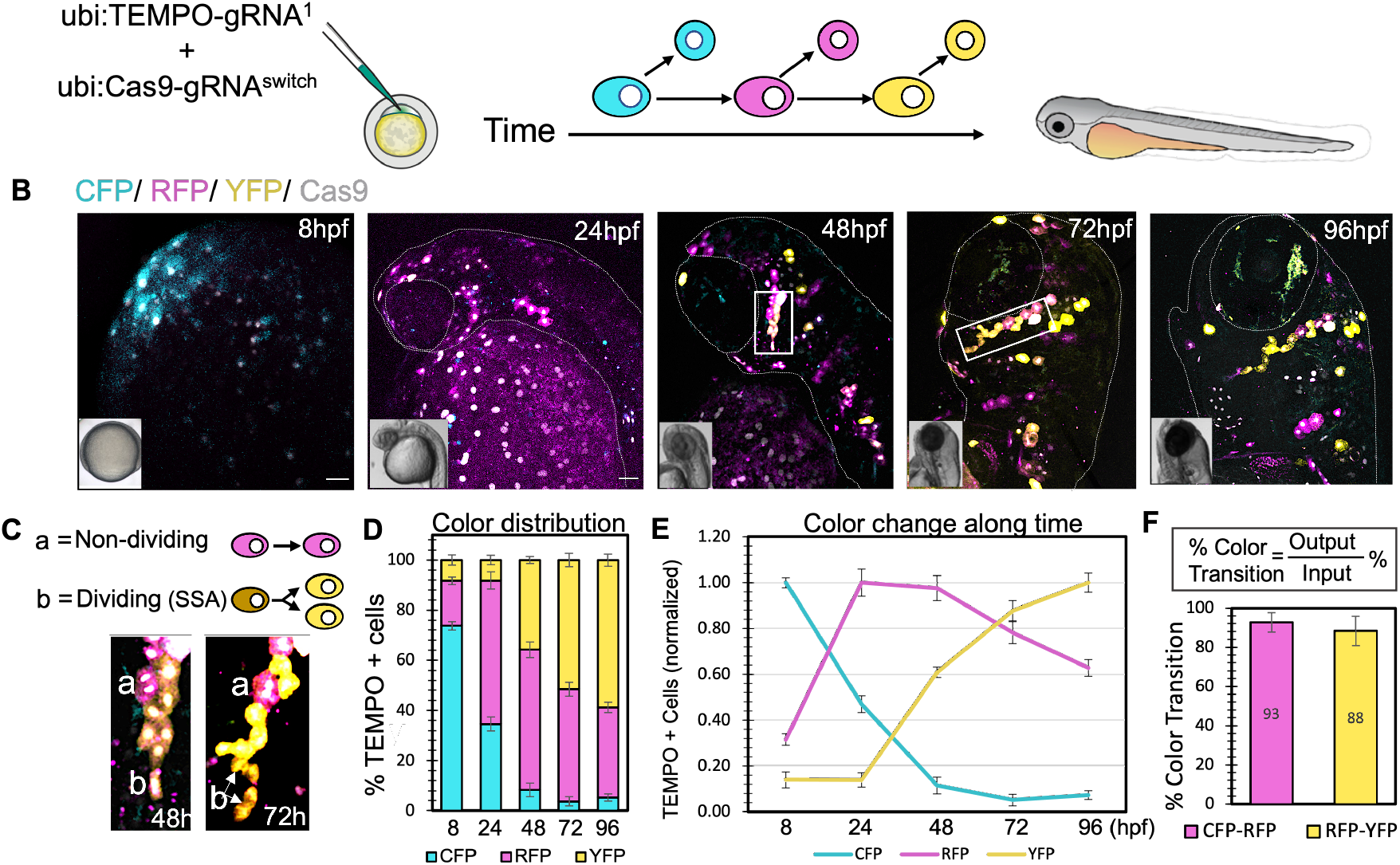
TEMPO works efficiently and enables sequential cell labelling in live zebrafish embryos. **(A)** Co-injection of Tol1 integrative TEMPO and Cas9 constructs into 1-cell-stage zebrafish embryos. TEMPO reporter transitions were analyzed in live embryos at subsequent developmental stages between 8 to 96 hours postfertilization. **(B)** Snapshots of the ubiquitously expressed TEMPO reporter cascade in live zebrafish embryos. Scale bar, 50 µm. **(C)** Insets highlight a cell clone imaged at 48 and 72h that contains examples of non-dividing cells (a) which do not transition in the cascade and dividing cells (b) which transition to the last color (YFP). **(D)** Changes in TEMPO color distribution (percentage of fluorescent cells for each color reporter along time) demonstrate efficient reporter transitions in a predefined order in developing zebrafish (8h, n=8; 24h, n=5; 48h, n=6; 72h, n=4; 96h, n=4, error bars represent SEM). **(E)** Normalization of the data shown in **(D)** to the maximum fluorescence for each reporter along time. **(F)** Efficiency of Color Transition, defined as the ratio between the final proportion of TEMPO+ cells at 96h (Output) with respect to the proportion of initial TEMPO+ cells at 8h (Input) for any given reporter transition. Data is shown as percentage ± SEM.

### TEMPO reveals spatio-temporal histories of progenitors and differentiated neurons in live zebrafish

To validate TEMPO in neuronal progenitors we focused on the atoh1a progenitor domain of the zebrafish embryonic hindbrain. This domain harbors dividing progenitors that differ in their cell mitotic index along the dorso-ventral and medio-lateral axis (*25*). For example, while progenitors located in the most dorsal part of the atoh1a domain do not divide or divide only once in a symmetric manner before differentiation, the adjacently ventral atoh1a progenitors can divide up to two times in a symmetric or an asymmetric manner. Further, dorsal atoh1a progenitors and derivatives migrate along the lateral-most part of the hindbrain, while ventral atoh1a cells migrate closer to the midline (Fig. 3A). To determine TEMPO dynamics in the atoh1a domain, we injected 1-cell-stage atoh1a:Gal4 embryos (*26*) with constructs expressing TEMPO downstream of a UAS promoter and Cas9 downstream of a her4.1 promoter, restricting Cas9 to neural progenitors and also co-expressing nuclear Halotag to identify Cas9+ cells (*27*) (Fig. 3B). Imaging of the zebrafish hindbrain at consecutive developmental stages and quantification of the distribution of TEMPO+ cells within different clones revealed that TEMPO progresses in a predetermined order in neuronal progenitors (Fig. 3C, D). We found most lateral atoh1a cells to be CFP+ or RFP+, indicating they do not divide or divide only once. However, YFP was consistently only expressed in medially located atoh1a progenitors at later stages, suggesting these cells divide enough to reach the last step of the reporter cascade (Figure 3E). These results are consistent with previous studies (*25*) and demonstrate that in addition to faithfully cell birth dating, TEMPO also predicts the dynamic history of cell proliferation and migration.

**Fig. 3.**
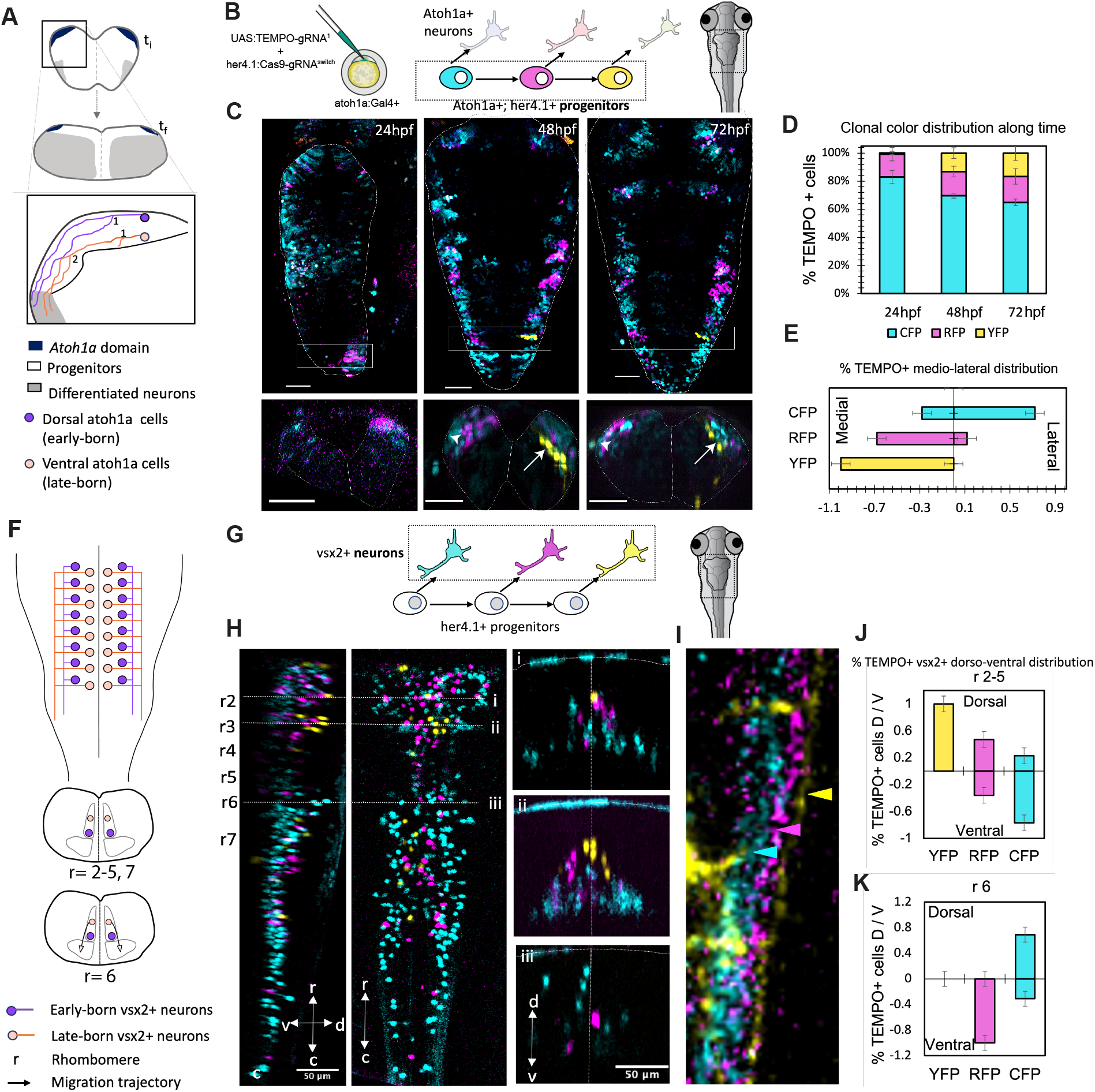
TEMPO reveals spatio-temporal cell histories of progenitors and mature neurons in the zebrafish hindbrain. **(A)** Schematic representation of the atoh1a progenitor domain in orthogonal hindbrain sections during early (ti) or late (tf) developmental stages. The inset in (A) highlights known differences in proliferation and migration dynamics between dorsal and ventral atoh1a progenitors (ref) (1, 2, first or second cell division, respectively). **(B)** To express the TEMPO fluorescent cascade in atoh1a progenitors of the zebrafish hindbrain, we injected two Tol1 constructs (*UAS:TEMPO-gRNA-1* and *her4.1:Cas9-gRNA-2 switch*) into 1-cell-stage zebrafish embryos of the transgenic line *atoh1a-Gal4*. **(C)** Representative images of consecutive TEMPO transitions in the atoh1a domain. Upper panels: Dorsal view of the zebrafish hindbrain; Lower panels: orthogonal z-projections of the insets in the upper panels. The arrow marks a representative YFP+ clone derived from a late-born progenitor and the arrowhead, a CFP+ clone derived from early-born progenitors. **(D)** Percentage of atoh1a fluorescent cells and color distribution along time. **(E)** The medio-lateral distribution of atoh1a derivatives correlates with TEMPO sequential reporter expression: early-born cells (CFP+) distribute laterally while later-born cells (YFP+) distribute medially. Cells born in-between these stages (RFP+) present an intermediate location which correlates with their birthdate. Data in (D) and (E) is shown as percentage ± SEM (n=5 fish, 3 independent experiments). **(F)** Cartoons represent the correlation between birthdate and spatial distribution of vsx2 neurons in dorsal (upper panel) and orthogonal (lower panels) views of the zebrafish hindbrain. **(G)** To activate the TEMPO reporter cascade in vsx2 neurons, we crossed a *vsx2:Gal4*;*UAS:TEMPO* transgenic line to a *her4.1:Cas9* line that expresses Cas9 in neuronal progenitors. **(H)** Lateral view (left) and dorsal view (center) of a stack from a representative fish expressing all transgenes at 6 days post-fertilization, showing a correlation between birthdate revealed by TEMPO sequential reporters and medio-lateral location. (Right) Orthogonal projections through the indicated rostro-caudal regions indicated in the left panels (i to iii) show a correlation between birthdate and dorso-ventral vsx2 distribution. **(I)** Detail of the distribution of TEMPO+ neuronal projections in a representative fish (color-coded arrows point to projections arranged medio-laterally by increasing birth-date). (J, K) Dorso-ventral distribution of TEMPO+ vsx2 neurons in rhombomeres R2-5 **(J)** or R6 **(K)**. Data shown as percentage ± SEM. n=5. Scale bars, 50 µm. hpf: hours post-fertilization.

In addition to provide information of neuronal progenitor dynamics, TEMPO can also be implemented to answer questions pertaining to spatial and temporal cell relationships of neuronal progenies. To this end we applied TEMPO to vsx2 interneurons, a group of excitatory neurons essential for locomotion (*28*). A previous study in zebrafish revealed that the birthdate of hindbrain vsx2 neurons determines their spatial distribution and function (*29*). While early-born vsx2 interneurons occupy lateral positions and produce rudimentary movements such as escape and struggle, late-born vsx2 interneurons are added medially and dorsally to the existing neurons and support more complex movements, including spontaneous slow swimming to search for food (Fig. 3F). To restrict TEMPO expression to this lineage, we crossed *UAS:TEMPO* to *vsx2:Gal4* fish (*28*). We then crossed *vsx2:Gal4; UAS:TEMPO to her4.1:Cas9* fish and determined the TEMPO color and spatio-temporal distribution of vsx2 interneurons in zebrafish larvae at 6 days-post-fertilization (dpf), when most neurons have ceased migration and differentiation (*29*). We found that early-born CFP+ neurons locate laterally to later-born RFP+ and YFP+ neurons, the latter being the population that locates closer to the midline (Fig. 3H). Orthogonal projections along different rostro-caudal levels of the hindbrain revealed early-born CFP+ neurons occupying ventral positions with respect to later-born RFP+ and YFP+ neurons (i and ii in Fig.3H). In contrast, at the level of rhombomere 6 in zebrafish, this pattern was inverted, where CFP+ neurons lied dorsally to later-born RFP+ (iii in Fig. 3H). This is consistent with previous studies showing that later-born neurons migrate ventrally in this hindbrain region, in contrast to other rhombomeres (*29*). Further, we demonstrated that early-born CFP+ neuronal projections lie medially to more lateral projections of RFP+ and the latest-born YFP+ neurons, as expected (Fig. 3I). These results demonstrate that TEMPO recapitulates known spatio-temporal patterns of differentiated neuron distribution in the same brain sample. Compared to previous approaches, TEMPO reveals previous cell division patterns and boosts the resolution of cell histories by increasing the number of sequential labels, while providing genetic access to specific temporal windows. Further, it allows specific spatial labelling when combined with the broadly used UAS/Gal4 system, making it a versatile strategy to reconstruct cell histories which could be applied to any tissue.

### TEMPO links birthdate of neurons and glia with layer distribution in the mouse brain

Existing technologies require live imaging for spatio-temporal reconstruction of cell histories (*8-9*). TEMPO circumvents this requisite, unlocking the potential of cell recording in other complex multicellular organisms with poor imaging accessibility such as mice. While different Cas9 modalities have been demonstrated to work in mouse (*30*), neither CaSSA reporters nor conditional gRNA switches have been tested before (*19-20*). To expand the applicability of TEMPO, we assessed its feasibility in mouse. We first modified our zebrafish constructs by replacing all promoters by ubiquitous promoters in mouse, generating two separated constructs (CAG-TEMPO-U6gRNA-1 and CAG-Cas9-U6gRNA-2switch) that integrate into the genome via PiggyBac (PB) transposition. We then tested the expression of the resulting constructs by in utero electroporation of embryonic mouse cortices (*31*) (Fig. 3A).

The mouse cortex offers an ideal system to perform spatio-temporal proof-of-principle experiments. Cortical neurons distribute in an ‘inside-out’ fashion: early-born neurons populate deeper neocortical layers (L6, then L5) while late-born neurons migrate past them to populate more superficial layers (L4, then layers L2-3) (*32*). To test TEMPO expression and cascade progression in cortical progenitors and neurons, we fixed mouse embryonic brains from E13.5 (24hours after the electroporation) to E17.5, after a 30-minute pulse of EdU to distinguish dividing progenitors. At E13.5, most cells were CFP+ progenitors located in the ventricular zone (VZ) and subventricular zone (SVZ) while no TEMPO+ cells were found in the cortical plate (CP). One day later, at E14.5, TEMPO progenitors had transitioned to RFP+, which located mostly to the VZ and SVZ, while CFP+ neurons were found migrating along the SVZ and into the CP. Only few YFP+ progenitors emerged from the VZ at this stage. The last TEMPO transition was most apparent at E15.5 when YFP+ progenitors appeared in the VZ while RFP+ neurons started to reach the CP.

Between E15.5 and E17.5, TEMPO+ progenitors emigrated from the VZ while later-born RFP+ (at E16.5) and YFP+ (at E17.5) neurons migrated past early-born CFP+ or RFP+ cortical plate neurons, respectively (Fig. 4B-D). To quantify the rate of color transition we focused on TEMPO progression in cortical progenitors. Sox2 expression was used to delineate the region occupied by radial glial cells (RGC) in the VZ (Fig.S4). Quantification of the percentage of color transition (% CT) resulted in 80% progenitor transitions from CFP+ to RFP+ in the first 24h after the electroporation. One day later, from E14.5 to E15.5, we found 31% progenitor transitions from CFP+ to RFP+ and 25% transitions from RFP+ to YFP+ (Fig. 4F, table inset). The lower transition capacity at later stages is consistent with the known decrease in cell division and reduction in the RGC numbers in the VZ during corticogenesis (*33*). This also explains the dramatic reduction in dividing TEMPO+ progenitors after E13.5 (Fig. 4F and S5). Overall, the progression of TEMPO in developing mouse cortices recapitulates the temporal dynamics of RGCs cell division capacity and sequential production of their progenies. Furthermore, these results demonstrate the efficacy of TEMPO to reconstruct spatio-temporal cell histories in a mammalian model.

**Fig. 4.**
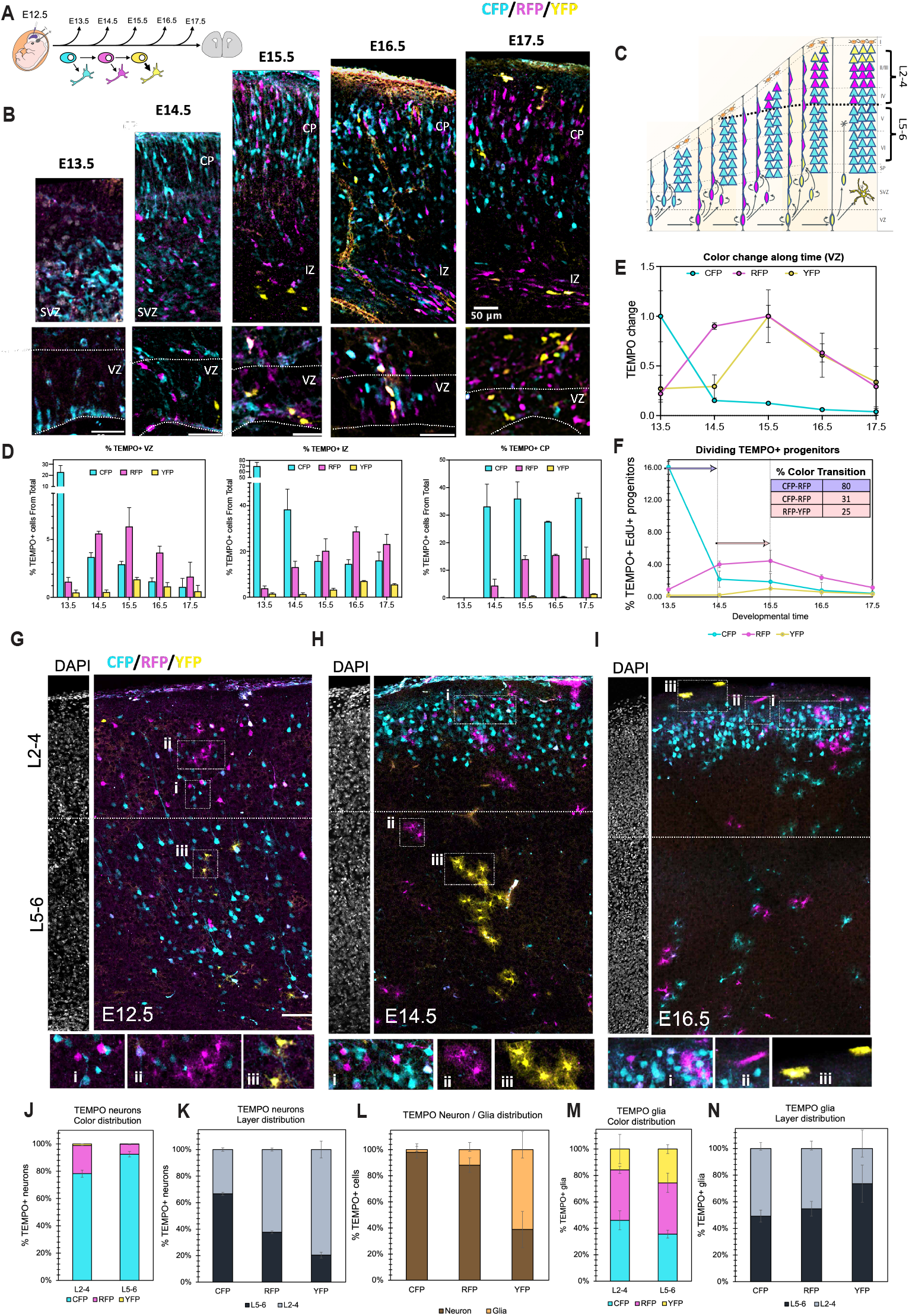
TEMPO connects sequential cell birth date and layer position in the mouse brain. **(A)** In utero electroporation (IUE) of TEMPO constructs into E12.5 mouse brains and analysis at subsequent developmental stages allow assessment of TEMPO reporter transitions during cortical development. **(B)** Progression of TEMPO cascade in neuronal progenitors follows a predefined order (CFP-RFP-YFP). Representative confocal images of mouse cortex at different stages following electroporation of TEMPO at E12.5. VZ, ventricular zone. SVZ, subventricular zone. IZ, intermediate zone. CP, cortical plate. Scale bars, 50 µm. **(C)** Schematic summary of the ‘inside-out’ distribution of sequentially labelled TEMPO neurons observed in the experimental samples. The X axis represents developmental time. **(D)** Distribution of TEMPO reporters in VZ, IZ and CP along time. Plots represent the mean percentage of fluorescent cells from the total of cells in that brain section SEM, 3 independent experiments. **(E)** Normalization of the VZ data shown in (D) to the maximum fluorescence for each reporter along time. Note that depletion of early CFP+ progenitors correlates with a rapid increase in RFP+ progenitors followed by an increase of late born YFP+ progenitors the following day. **(F)** Proliferative capacity of neural progenitors labelled with TEMPO reporters decreases along time. In correlation, the efficiency of color transition is higher at earlier stages (E13.5 to E14.5, purple arrow and corresponding table row) than at later stages (E14.5 to E15.5, pink arrow and corresponding table rows). (G-I) Postnatal P10 brain sections from mice electroporated with TEMPO constructs at stages E12.5-E16.5. DAPI labelling (shown as separated adjacent panels) was used to determine the limit between upper (L2-4) and lower (L5-6) cortical layers (dashed line). Insets highlight representative examples of TEMPO+ neurons **(i)** and astrocytes (ii-iii). **(J)** Percentage of neurons expressing each TEMPO reporter in the upper (L2-4) or lower (L5-6) layers. **(K)** Layer distribution of TEMPO+ neurons for each reporter reveals an ‘inside out’ pattern. Early CFP+ neurons mostly occupy lower layers while later born RFP and YFP neurons occupy upper layers. **(L)** Neuron and glia ratio for each reporter. **(M)** Percentage of astrocytes expressing each TEMPO reporter in the upper (L2-4) or lower (L5-6) layers shows that most astrocytes are labelled by later reporters (RFP+ and YFP+) in both compartments. **(N)** Layer distribution of TEMPO+ astrocytes for each reporter does not show a significant correlation between astrocyte birthdate and layer distribution. Data is represented as percentage SEM. n=3, Scale bar (G-I), 100 µm.

We then sought to explore TEMPO color distribution in postnatal mouse brains, 10 days after birth (P10), when cortical layer formation and neuronal migration are complete and astrocytes can be distinguished morphologically (*32*). Electroporation of TEMPO constructs at different developmental stages allowed temporal subdivision of corticogenesis starting from either the generation of deep layers (E12.5) or upper layers and gliogenesis (E14.5 and E16.5, respectively), following the expected inside-out pattern of neuronal lamination (Fig. 4G-I). These results also demonstrated that cascade progression works independently of the stage of the targeted progenitors. Electroporation at E12.5 and analysis of P10 brain cortices showed that CFP+ neurons occupy mostly the lower layers (L5-6) consistent with an early colonization, while later-born RFP+ and YFP+ neurons occupy mostly the upper layers (L2-4) (Fig. 4J, K). The proportion of neurons labelled with later colors (RFP+ and later YFP+) was lower than that for the first color (CFP+) consistent with lengthening of the cell cycle and the decrease in cell division (Fig. 4J) (*33*). While overall distribution of TEMPO+ neurons was as expected from the inside-out pattern of layer formation, we found a high proportion of CFP+ neurons in the upper layers (Fig. 4J, K). This could be attributed to the postmitotic status of a high proportion of CFP+ cells immediately after electroporation, which would impair cascade transitioning. To test this hypothesis, we co-electroporated an episomal control plasmid (CAG-Halotag) and the TEMPO integrative constructs at E12.5. The episomal plasmid is diluted with each cell division while TEMPO constructs integrate into the genome and propagate throughout development. We found that more than 28% CFP+ neurons in the upper layers co-expressed the episomal plasmid, and thus did not divide much after the electroporation. In contrast, less than 10% of RFP+ neurons and none of the YFP+ neurons co-expressed the episomal plasmid, consistent with their progenitors undergoing more divisions before differentiation (Fig. S6).

After neurogenesis is complete (∼E16), neural progenitors transition to a gliogenic mode, generating astrocytes and oligodendrocytes (*32*). Given the timing of the transition from neuro to gliogenesis and that of TEMPO reporters electroporated at E12.5 (Fig. 4B), we reasoned that the proportion of cells labelled with each TEMPO color that differentiate into glia should increase with cascade progression. The quantification of TEMPO+ astrocytes distinguished by morphology revealed that while most CFP+ and RFP+ cells were neurons, most YFP+ cells were astrocytes (Fig. 4L). We still found some CFP+ astrocytes but this was explained by their lack of Cas9 expression, which would not allow cascade progression (Fig. S7). In contrast to TEMPO+ neurons whose distribution was highly correlated with the inside-out pattern of layer formation (Fig. 4B, G, K), TEMPO+ astrocytes were scattered along the radial axis (Fig. 4G-I, N), consistent with previous studies showing that astrocyte spatial localization is highly plastic (*34*). Interestingly, analysis of TEMPO color distribution in superficial Layer 1 (L1) astrocytes revealed a dramatic reduction in cascade progression in brains electroporated at E16.5 compared to brains electroporated at earlier stages (E12.5-E14.5), suggesting most L1 astrocytes are generated perinatally and do not divide much after that (Fig. S8).

Overall, these results demonstrate that TEMPO works in a predefined order and recapitulates the inside-out pattern of cortical layer formation and the transition from neurogenesis to gliogenesis in mouse.

### TEMPO-2.0 allows sequential genetic manipulation in mouse cortical progenitors

Cell specification during tissue and organ formation often relies on temporal patterning: different cell types are produced sequentially when exposed to temporal intrinsic or extrinsic cues. An evolutionary conserved strategy for cell specification relies on deploying cascades of transcription factors in a particular order (*4-6*). Determining the mechanisms of cell type specification and being able to expand specific cell-types at will, would require tools capable of genetic manipulation in a predefined order, enabling sequential activation and inactivation (temporal “pulses”) of effector genes in specific developmental windows. Such tools are not available in vertebrates and we envision TEMPO to overcome that limitation. To this end, the modular design of TEMPO allowed us to express effector genes in different temporal windows by incorporating the effector sequence in frame with the correspondent reporter gene (Fig. 5A-B). Thus, TEMPO-2.0 couples activation and inactivation of a reporter in a particular temporal window with that of an effector gene, leaving the other reporter frames intact and allowing sequential labelling of manipulated and control cell populations.

**Figure 5.**
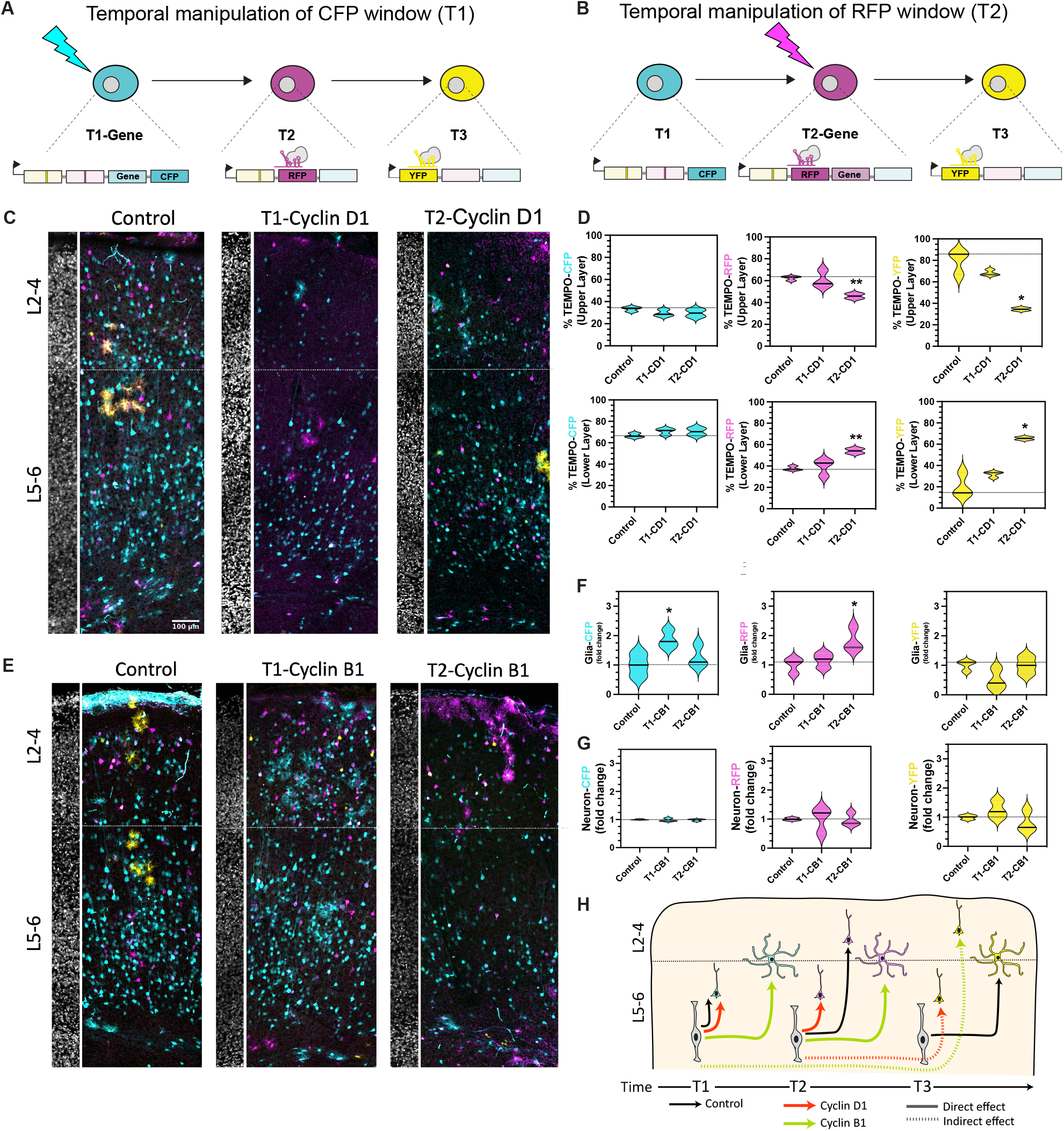
Temporal genetic manipulation of cell cycle regulators in mouse brain progenitors. **(A, B)** Design of TEMPO-2.0 constructs overexpressing a transgene in the first temporal window with the CFP+ reporter (A) or in the second temporal window with the RFP reporter **(B). (C)** Representative P10 sections through the cortex of control (left panel) or overexpressing Cyclin D1 in the first (T1) or second (T2) temporal window. DAPI labelling establishes the limit (dash line) between upper (L2-4) and lower layers (L5-6). **(D)** Violin plots represent layer distribution of neurons labelled by each color reporter in control and manipulated neurons. CD1, Cyclin D1. n=3. Scale bar, 100 µm. We found a significant shift in layer distribution for RFP+ (^**^p<0.01) and YFP+ (^*^p<0.05) neurons from upper to lower layers after Cyclin D1 overexpression in the second temporal window (a two-tailed unpaired Student’s t test was used). **(E)** P10 cortical sections of control (left panel) or overexpressing Cyclin B1 in the first (T1) or second (T2) temporal window. (F, G) Violin plots represent the change in astrocyte numbers **(F)** or neuron numbers **(G)** (normalized to the maximum for each color reporter) in control or manipulated samples. CB1, Cyclin B1. n=3. Scale bar, 100 µm. We found a significant increase in astrocytes labelled CFP+ or RFP+ when overexpressing Cyclin B1 in the first (T1) or second (T2) temporal windows, respectively (^*^p<0.05) (a two-tailed unpaired Student’s t test was used). **(H)** Summary of results obtained after TEMPO or TEMPO-2.0 electroporation. Schematic representation of the mouse cortex divided in upper (L2-4) and lower (L5-6) layers. Apical progenitors (grey elongated cells) generate neurons that colonize the cortex in an ‘inside-out’ fashion. In control samples (black arrows), early-born CFP+ neurons occupy lower layers, later-born RFP+ neurons mostly occupy upper layers and the latest-born YFP+ occupy upper layers and mostly produce astrocytes found scattered in the cortex. Overexpression of Cyclin D1 (orange arrows) shifts neuron distribution to lower layers in the corresponding temporal window. Overexpression of Cyclin B1 (green arrows) causes a shift towards late cell fates and increases the number of astrocytes in the corresponding temporal window. Indirect effects on other non-perturbed temporal windows, such as the increase in late-born YFP+ neurons by overexpression of Cyclin B1 in the first temporal window, are depicted by dashed lines.

In multiple developmental contexts, cell cycle and cell fate decisions are strongly linked. For example, longer cell cycles due to a longer G1-phase have been associated with fate-decision making (*35*). Interestingly, several components of the cell cycle machinery are not uniformly expressed throughout neurogenesis suggesting a role in neuronal type temporal regulation (*36*). Recent scRNA-Seq studies showed that early cortical progenitors express higher levels of Ccnd-1 (Cyclin D1-required during G1/S) than later progenitors while Ccnb-1 (Cyclin B1 – required during G2/M) expression is maintained in later progenitors (*37, 33*). The temporal change in Cyclin D1 expression levels coincides with the transition from lower to upper layer neurogenesis and suggests a link between Cyclin D1 expression and the generation of early-born-lower layer neurons.

To test this hypothesis and demonstrate we could manipulate the spatio-temporal distribution of mouse cortical progenies with TEMPO-2.0, we incorporated sequences of either Cyclin D1 or Cyclin B1 to be expressed in different temporal windows (Fig. 5A, B). We then co-electroporated the resulting plasmid with a Cas9 containing plasmid at E12.5 and analyzed postnatal P10 mouse brains. As a control, we used the original TEMPO construct, in which no perturbation was introduced. A pulse of Cyclin D1 in the first temporal window (T1-Cyclin D1) slightly increased the number of CFP+ lower layer neurons at the expense of CFP+ upper layer neurons. This tendency was also observed in RFP+ and YFP+ neurons (Fig. 5C, D). A pulse of Cyclin D1 in the second temporal window (T2-Cyclin D1), showed no major effect on CFP+ neurons with respect to the control, but revealed significant changes in the distribution of RFP+ and YFP+ neurons, which concentrated in the lower layers instead of the upper layers (Fig. 5C, D). Our results strongly suggest that Cyclin D1 overexpression shifts lineage commitment towards lower layer neurons in the manipulated temporal window, a phenotype especially evident in the second temporal window (T2∼ E14.5-15.5), when Cyclin D1 expression would have been normally reduced (33). Furthermore, we show that perturbation of specific temporal windows also impacts cells in subsequent temporal windows, while leaving the preceding lineages intact. These findings are in contrast with previous work which concluded that Cyclin D1 overexpression forces cortical neuron differentiation towards the cell-fates normally generated at the time of induction, rather than promoting a specific lower layer neuronal cell fate. Previous approaches, however, did not allow simultaneous visualization of manipulated and control neurons born at different developmental times, which is now possible with TEMPO-2.0.

Contrary to Cyclin D1, Cyclin B1 has been linked with progenitor maintenance and later-born cortical cell generation (*37*). We thus analyzed the effects of temporal overexpression of Cyclin B1 with TEMPO-2.0, reasoning that this could shift perturbed temporal windows to later lineages, including astrocytes. We found that overexpression of Cyclin B1 in the first temporal window (T1-Cyclin B1) increased the number of CFP+ astrocytes but not that of RFP+ astrocytes in the subsequent window. In contrast, overexpression of Cyclin B1 in the second temporal window (T2-Cyclin B1) increased RFP+ astrocytes, while CFP+ astrocyte numbers remained comparable to control samples (Fig. 5E, F). These results demonstrate that Cyclin B1 overexpression leads to a shift from neurogenesis to gliogenesis in the specific temporal window where it is activated (T1 or T2). Interestingly, early Cyclin B1 overexpression (T1-Cyclin B1) caused a slight increase in late-born YFP+ neurons which could be explained by an increase in progenitor maintenance enhancing cell division capacity, thus providing more opportunities for reporter cascade progression within the neurogenic period. In parallel, we observed a decrease in YFP+ astrocytes, which suggests an early YFP progenitor pool exhaustion during the neurogenic phase (Fig. 5F, G).

Taken together, these results demonstrate that TEMPO-2.0 can be used to sequentially label and manipulate neural progenitors within predefined temporal windows and simultaneously analyze the phenotype of perturbed and control neurons and glia (Fig. 5H). This enables the study of temporal effects of molecular factors on cell specification.

## DISCUSSION

Establishing the association between a cell’s birth timing and tissue distribution is essential to define mechanisms of cell specification, not only in development but also during regeneration and in disease states. However, the dynamic nature of these processes has made it challenging to address in most complex organisms. TEMPO overcomes this limitation by labelling successive cell generations with different colors, revealing temporal and spatial cell relationships in single vertebrate samples, including zebrafish and mouse. Our most recent version, TEMPO-2.0, further allows temporal manipulation of regulatory factors in cycling cells in a predefined order. This permits simultaneous analysis of perturbed and control progenies, enabling functional studies of serial temporal fates in the same tissue.

Contemporary approaches for cell lineage tracing use cumulative editing of DNA recorders and rely on non-ordered edits to retrospectively reconstruct cell histories (*15*). One big limitation of these strategies is that similar outcomes could result from different editing orders or from large deletions which erase previous edits, confounding temporal reconstruction (*16-17*). Proof of concept experiments in live zebrafish embryos revealed that TEMPO overcomes these limitations by introducing reporter edits in a predefined and invariant sequence. There is no other available strategy in vertebrates that allows assessment of the status of a lineage recorder along time in the same living sample, given that other readouts involve tissue fixation or dissociation (*9*). In the future, TEMPO could be combined with barcoding methods to drive recorder editing in a predefined order while extending recorder availability over longer periods of time (*17*). Further, combining TEMPO with high-density clonal labelling methods such as intMEMOIR (*38*), or Brainbow (*39*), to distinguish cell clones while maintaining both temporal and spatial resolution would be very powerful for multiplexed lineage tracing in complex organisms. In addition, TEMPO is compatible with spatial-transcriptomic methods (*40*) enabling combined studies of cell identity and temporal patterning.

The versatile design of TEMPO permits combination with available transgenic lines for conditional spatio-temporal control of expression (Gal4/UAS, tTA/TRE, heat shock). The transgenic zebrafish line her4.1:Cas9 allows Cas9 expression in most neural progenitors and could be combined with UAS or TRE-based TEMPO lines and available Gal4 or tTA lines for neuron-specific analysis. Targeting TEMPO to known zebrafish neuronal lineages enabled reconstruction of spatio-temporal relationships of different sublineages in single samples (Fig. 3). This is in stark contrast with the high number of samples required to obtain the same conclusions with pre-existing techniques (*25, 29*). In addition to studying neuronal lineages, the conditional TEMPO zebrafish lines can be combined with other drivers to enable temporal labelling of any tissue of interest that contains dividing cells.

Temporal access to longer lineages using TEMPO could be useful but would require adding steps to the color cascade. We proved that new gRNA sequences worked robustly and had a similar transition efficiency (>70%) to the one used in this study (Fig. S1, S2) so that adding more steps is feasible. Modifying cascade speed may be desired as well. To that end, coupling Cas9 expression to the overexpression of enzymes in charge of SSA repair may help shift the repair mechanism towards SSA instead of non-homologous end joining (NHEJ) or other competing repair mechanisms (*21*).

A major advantage of TEMPO is that it allows spatio-temporal reconstruction of cell histories retrospectively in single samples without the need to perform live imaging. This is an essential feature to be able to access cell histories in complex organisms with poor imaging accessibility, such as the mouse. We thus implemented TEMPO in mouse, by targeting the transgenes to developing mouse brains by in utero electroporation. This allowed, for the first time, to subdivide cortical lineages in three consecutive developmental windows (separated from each other by around 24hours) recapitulating the inside-out pattern of neuronal layer formation and the early-to-later shift from neuro to gliogenesis in single cortex samples (Fig.4). Currently, only MADM (Mosaic Analysis with Double Markers) (41) can access mouse cortical lineages with single-cell resolution. This technology is based on rare inter-chromosomal recombination events which label two daughter cells with different colors, allowing simultaneous labeling and gene knockout in single cells. However, MADM requires many samples to obtain general conclusions on cell lineage and does not allow compared studies between temporally distinct populations. By contrast, TEMPO labelling and manipulation of subsequent cell populations allows a dramatic reduction in the number of samples required to map full sublineages, which will speed up biological discovery. By allowing simultaneous visualization of cells born at different times, TEMPO reveals important differences in cell dynamics which would otherwise be overlooked without temporal resolution. For instance, previous studies predicted that direct progeny of apical progenitors colonize the superficial layers early in development but could not provide definitive proof because of the lack of temporally distinct labels for later born neurons (*42*). Our observations that a significant proportion of early born neurons (labelled by the first TEMPO color (CFP) and an episomal plasmid (Fig. S6)), colonize the upper cortical layers provide further proof that direct progeny of apical progenitors colonize the superficial layers.

Previous studies using multicolor clonal labelling concluded that astrocytes, which are produced perinatally, colonize the cortex in a non-ordered fashion where a single progenitor can produce superficial and cortical parenchyma astrocytes (*34*). However, those studies used clonal labelling techniques which lack temporal resolution and cannot distinguish subsequent generations of astrocytes in the same sample. By electroporating TEMPO constructs at different stages, one can subdivide different developmental processes of interest into temporal windows. As an example, electroporation of TEMPO constructs into E16.5 mouse brains resulted mostly in TEMPO+ astrocyte labelling and only few CFP+ neurons were labelled. This result was expected, given neurogenesis is almost complete at this stage and gliogenesis is underway. Interestingly, TEMPO electroporation at consecutive developmental stages showed a decrease in reporter cascade progression in superficial L1 astrocytes precursors along time (Fig. S8). This suggests that most superficial L1 astrocytes are generated perinatally and do not divide much after that, linking proliferation mode and cell-fate. This further highlights the importance of TEMPO in revealing otherwise unnoticed cellular spatio-temporal relationships.

Beyond spatio-temporal cell labelling, the design of TEMPO enables the activation of genetic cascades, ideal for functional analysis in vivo. We demonstrated that TEMPO-2.0 allows perturbation and differential labelling of temporal windows in a precise order and visualization of both control and perturbed progenies in the same sample (Fig. 5). Overexpression of Cyclin B1 or Cyclin D1 in different temporal windows resulted in a fate switch from neurons to glia or a change in neuronal layer distribution from upper to lower layers of the perturbed progeny, respectively (Fig. 5). Interestingly, we show that shifting the commitment of early progenitors to later fates (astrocytes) through Cyclin B1 overexpression, seems to exclusively affect those progenies arising from the perturbed window, while later-born progenitors remain capable of producing neurons and have a tendency towards enhanced neurogenesis (Fig. 5E-H, S9). This suggests a feedback regulation which could be compensating the early generation of astrocytes at the expense of neurons.

Previous studies suggested that Cyclin D1 overexpression promotes cortical neuron differentiation towards cell-fates normally generated at the time of induction. By contrast, our results suggest that Cyclin D1 overexpression promotes a specific temporal fate, shifting neuronal commitment towards lower layer neurons not only in the manipulated temporal window but also in the following windows. A potential explanation for this discrepancy is that previous cell labelling approaches relied on electroporation of episomal plasmids, which only label post-mitotic cells that are few cell cycles away from the electroporated radial glial progenitors. Given that cell cycles of radial glial progenitors are not synchronized (44) and that the exact number of divisions labelled by episomal plasmids is unclear, it is difficult to estimate the effect of manipulating cell cycle regulators through episomal plasmid labelling. By contrast, TEMPO tracks consecutive cell generations in the same sample by incorporating stable labels, crucial for studying serial temporal fates in cycling neural stem cells. Our experiments, showing a persistent effect of Cyclin D1 for the neuronal commitment towards lower layers, suggest that cortical progenitors are susceptible to the action of Cyclin D1 along an ample window of developmental stages. Therefore, TEMPO 2.0 unlocks the potential to explore whether late cortical precursors are still permissive to temporal fate manipulation and determine when cell-fate becomes irreversible.

In sum, TEMPO is a broadly applicable imaging-based system for temporal genetic labelling and manipulation. Its compact and versatile design dependent on CRISPR/Cas9 makes it compatible with most in vitro and in vivo models, and possible to combine with available transgenic lines to refine spatio-temporal targeting. TEMPO-2.0 enables modification of the cell specification cascade by introducing effectors at predefined temporal windows and could be used to define the temporal boundaries in which differentiation factors can regulate cell fates. TEMPO could not only improve our understanding of the temporal mechanisms regulating cell specification but could also be used to provide a predefined sequence of factors to instruct the generation of cell subtypes. Such cellular engineering could revolutionize in vitro disease modeling and stem-cell therapy applications.

## Acknowledgments

We thank all members of Tzumin’s lab for their comments and feedback. We thank Claude Desplan and Erik Snapp for critical reading and revision of the manuscript. We also thank Sarah Lindo, Kendra Morris, and the Janelia histology and vivarium core facilities for their excellent technical support. We thank Kathryn Miller for administrative support;

## Author contributions

Conceptualization, I.E.-M., D.F and T.L.; Methodology, I.E.-M. and T.L. Investigation, I.E.-M., D.F, C.B.-M.; Writing – Original Draft: I.E.-M; Writing – Review & Editing, I.E.-M., D.F, R.L.M., J.G.-M., C.B.-M., C.P., M.K. and T.L.; Visualization, I.E.-M, Supervision, M.K. and T.L.; Project Administration, M.L. and T.L.; Funding Acquisition, M.K. and T.L.;

## Data and materials availability

all plasmids generated in this study will be available through Addgene after publication and the zebrafish transgenic lines will be made available through the terms of a Material Transfer Agreement (MTA).

## Supplementary Materials

## Material and Methods

### Plasmids cloning

DNA constructs were designed using Benchling (https://benchling.com) and cloned using standard methods, including PCR, restriction digest, Gibson assembly, Gateway cloning and verified using Sanger sequencing. Constructs containing repeated regions (gRNA switches and TEMPO reporters) were cloned combining multiple synthetic DNA blocks or PCRs by Gibson assembly. TEMPO reporters in their inactive state contain 100bp long repeats flanking a target gRNA sequence. The improved version of the inactive gRNA switch used in this study (Fig. S1) contains 43bp long repeats flanking a target gRNA sequence. Details for each construct and gRNAs reported in this manuscript can be found in Table S1.

### Zebrafish injections

Zebrafish adults (3 months-2 years old) were mated to generate embryos. Tol2 and Tol1 mRNA were synthetized from linearized DNA using the mMessage mMachine SP6 Transcription kit (Thermo Fisher 995 Scientific) and purified (RNAeasy Mini Kit, Qiagen) before injection. About 200 embryos for each experiment were injected at 1-cell-state with 1-2 nanoliters of 25 ng/ul of Tol2 or Tol1 transposase mRNA and 25 ng/ul of the corresponding Tol2 or Tol1 integrative plasmid. When several plasmids were co-injected, the total amount of DNA did not exceed 40ng/ul.

### Zebrafish transgenic lines

The following transgenic lines were used in this study: *atoh1a:Gal4* (*26*), *vsx2:Gal4* (*28*). We generated *UAS:TEMPO_gRNA-1* and *her4.1:Cas9 H2B-Halotag_gRNA-2switch* transgenic lines by Tol transposition (see Zebrafish injections above). Embryos used for these injections include: *vsx2:Gal4* embryos for *UAS:TEMPO_gRNA-1* injections to obtain double transgenic zebrafish, and Casper embryos for *her4.1:Cas9-H2B-Halotag_gRNA-2switch* injections. Screening of *vsx2:Gal4:UAS:TEMPO* positive founders and subsequent generations was based on correct pattern of expression and coverage of the *vsx2* domain (*28-29*). Expression of Halotag protein along with Cas9 in *her4.1:Cas9-H2B-Halotag_gRNA-2* zebrafish was used to screen positive founders and their progeny. We sorted Halotag positive zebrafish by pre-loading 30 nM Halotag dye JF568 in 5mL of zebrafish water two hours prior to screening on a fluorescent binocular, using the RFP channel. Bright nuclear Halotag expression in the *her4.1* neural progenitor domain (*27*) was used to select zebrafish for propagation.

### Mouse *in utero* electroporation

*In utero* electroporations were performed in embryonic day E12.5, E14.5 or E16.5 timed-pregnant C57BL/6J mice (Charles River). Mice were anesthetized by using an isoflurane-oxygen mixture [2% (vol/vol) isoflurane in O2]. The uterine horns were exposed and 1 μL of DNA solution was pressure-injected through a pulled glass capillary tube into the right lateral ventricle of each embryo. The solution contained a mixture of *CAG:TEMPO_gRNA-1* and *CAG:Cas9_gRNA-2* integrative plasmids and a *CAG:PiggyBac* transposase expressing plasmid at a 1:1:1 molar ratio, with a maximum concentration of 2 μg/μL. Immediately after injection, the head of each embryo was placed between tweezer electrodes and four pulses of 50 ms and 100 V were applied at 950 ms intervals. Electroporated embryos at day E13.5 to E17.5 were dissected and fixed overnight in 4% PFA and postnatal mice were perfused with cold saline and 4% PFA and postfixed overnight.

All animal experiments were conducted according to the National Institutes of Health guidelines for animal research. Procedures and protocols (19-179) on mice were approved by the Institutional Animal Care and Use Committee at Janelia Research Campus, Howard Hughes Medical Institute. Animals were kept with a 12 h dark/12 h light cycle in a temperature controlled (20–22 °C, humidity: 30–70%) and sound attenuated room.

### Histology

Mice brains at early embryonic stages (E13.5-E15.5) were embedded in a 15% Sucrose and 7.5% Gelatin mix solution, frozen at -80°c and sectioned (25 μm) using a Leica Cryostat. Primary antibody staining was performed overnight on sections in a 1xPBS, 0.1%Triton X-100, 20%FCS (fetal calf serum) solution. The slides were washed x3 times in PBS-0.1%Triton X-100 and secondary staining was performed in 1xPBS, 0.1%Triton X-100, 2-4 hours at room temperature. Slides were mounted with Fluoromount (Sigma, Cat. # F4680) and analyzed with a Zeiss LSM880 confocal microscope. Late embryonic and postnatal mice brains (E16.5-P10) were embedded in a 5% agarose-PBS solution and sectioned (100 μm) using a Leica Vibratome. Staining was performed in blocking buffer (PBS, 2% BSA, 0.1% Triton X-100). Primary antibody incubation was performed overnight at 4C. Sections were washed x3 times in blocking buffer for a total of 45 minutes and incubated with secondary antibodies for 2-4 hours at room temperature. Sections were mounted on slides with Fluoromount and analyzed with a Zeiss LSM880 confocal microscope. The following primary antibodies were used: mouse anti-V5-tag to amplify V5-tagged CFP (Thermo Fisher, R96025, 1:650), rat anti-mCherry (Thermo Fisher, M11217, 1:500), rabbit anti-Halotag (Promega, G9281, 1:500), rabbit anti-Sox2 (Abcam, ab97959, 1:1000). The following secondary antibodies were used: Alexa Fluor goat anti-mouse 647 (Thermo Fisher, A-21235, 1:500) or goat anti-mouse 405 (Jackson Immunoresearch, 715-476-150, 1:500), Alexa Fluor goat anti-rat 568 (Thermo Fisher, A-11077, 1:500) and Alexa Fluor goat anti-rabbit 647 (Thermo Fisher, A-21244, 1:500).

### EdU labelling

To identify dividing TEMPO positive progenitors in the ventricular zone of developing mouse embryos at the time of the analysis (E13.5-E17.5), 50 mg/kg body weight of a 8 mg/ml EdU (Thermo Fisher Scientific) stock solution was administered to pregnant mice by intraperitoneal injection 30 min prior to embryo dissection. EdU was detected using the Click-iT EdU Alexa Fluor imaging kit (Thermo Fisher Scientific, C10340, 647 staining), according to manufacturer’s protocol. Briefly, cortex cryostat sections or floating vibratome sections were first incubated with primary and secondary antibodies and after 3x washes in PBS, sections were incubated for 30 mins with the Click-iT reaction cocktail, protected from light. Sections were washed several times in PBS before mounting using Fluoromount (Sigma).

### Image acquisition and processing

Live zebrafish embryos and larvae were first anesthetized by bath application of 0.02% w/v solution of 1025 Ethyl-3-aminobenzoate methanesulfonate (Sigma-Aldrich, St. Louis) in filtered fish system water for 1 min. Fish were then mounted in a drop of 0.8% low melting point agarose (Invitrogen) over a glass-bottomed plate and imaged using an inverted Zeiss LSM 880 confocal microscope. For timed snapshots of live zebrafish along development we removed mounted embryos after each imaging session and placed them in fresh water until the next session.

Mice brain sections were imaged on a Zeiss LSM 880 confocal microscope and Fiji (NIH) was used for downstream image processing and analyses.

### Quantitative analysis

#### Zebrafish distribution of TEMPO clones

We calculated the medio-lateral distribution of TEMPO positive *atoh1a* progenitors as follows: we first established the center of each selected clone in Fiji and quantified the number of cells expressing each TEMPO reporter at a given distance to that reference point. Cells closer to the hindbrain midline were considered medially located and cells closer to the surface were considered laterally located.

#### Mice quantitative analysis

The total number of TEMPO+ cells expressing each color reporter in embryonic or postnatal mice brains was recorded using FIJI Cell Counter. Regions of interest (ROI) including the ventricular zone (identified by Sox2 labelling during embryonic stages), and the lower and upper cortical layers (identified using DAPI staining) were determined prior to the analysis. Statistical significance calculations comparing two conditions (control and perturbed temporal window in Fig.5 and Fig. S9) were performed using a two-tailed unpaired Student’s t test. Results are expressed as mean ±SEM %.

---

**Fig. S1.**
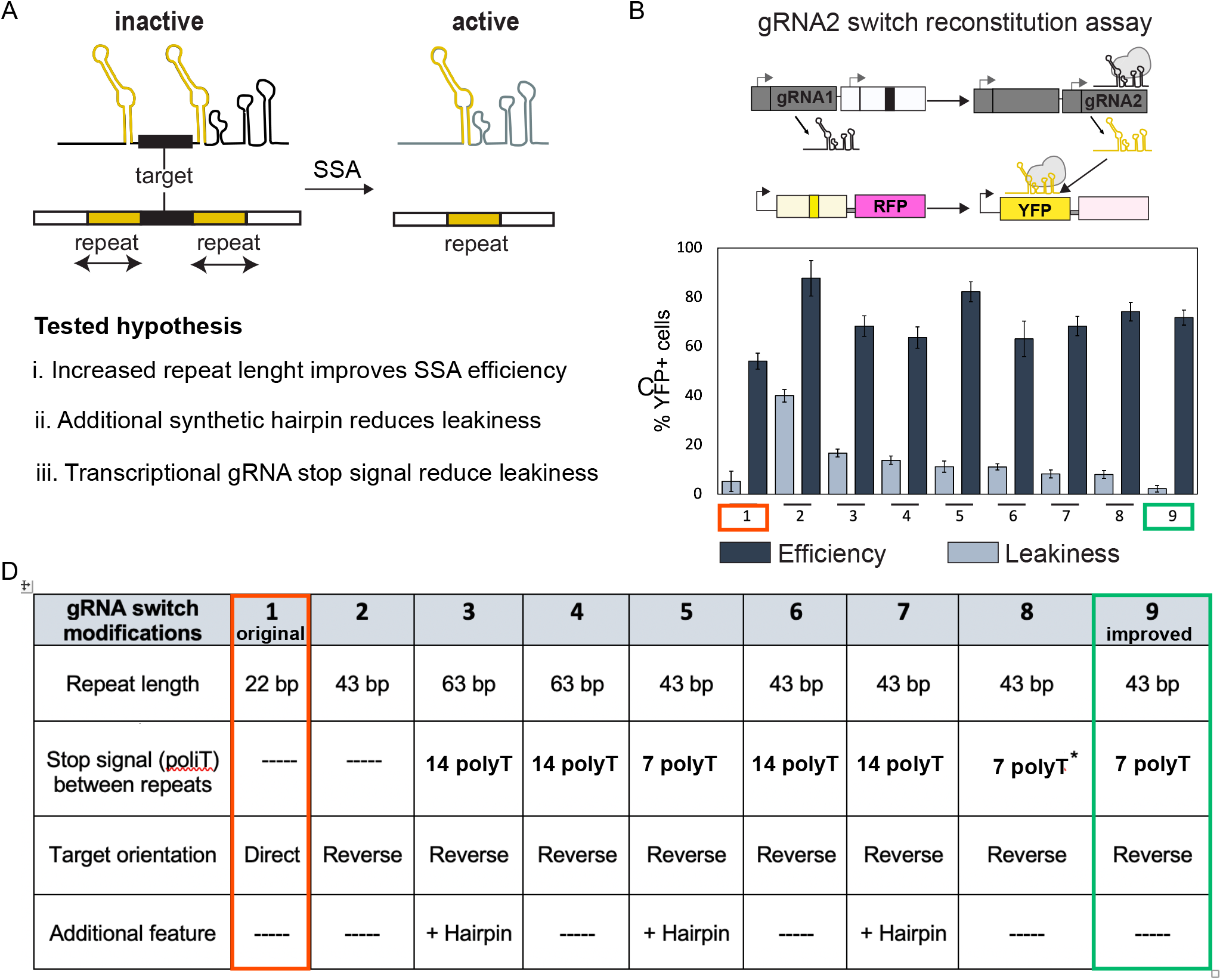
Improving gRNA switch repair efficiency. (**A**) gRNA switch repair scheme. Repeats located in the scaffold region are highlighted in yellow. We tested whether increases in repeat length improve repair efficiency (i); or incorporation of a new hairpin motif, adjacent to the target sequence, reduces leakiness of the inactive gRNA switch by changing its secondary structure; or whether integration of a polymerase-III termination signal (polyT) reduces leakiness of the inactive gRNA switch (*45*). (**B**) gRNA switch fluorescent assay to test gRNA-2 switch repair. In the presence of gRNA-1 and Cas9, gRNA-2 is activated and consequently, YFP is repaired. (**C**) Efficiency and leakiness of the 9 tested gRNA switch variants, assessed by expression of YFP reporter in 3 day-post-fertilization (dpf) zebrafish larvae injected at 1-cell-stage (mean ± SEM; total n=116 fish, per condition n= 7-22). The original gRNA switch (*19*) and improved variant chosen are highlighted in red and green, respectively. (**D**) Table legend of characteristic features for each variant tested in this study. The polyT termination signal was placed downstream of the target sequence between the scaffold repeats in the inactive gRNA, except for variant 8, in which the polyT was placed upstream (asterisk). The target orientation is determined by the location of the PAM (protospacer adjacent motif) sequence and it is downstream of the target sequence in the Direct orientation, while it is upstream in the Reverse orientation.

**Fig S2.**
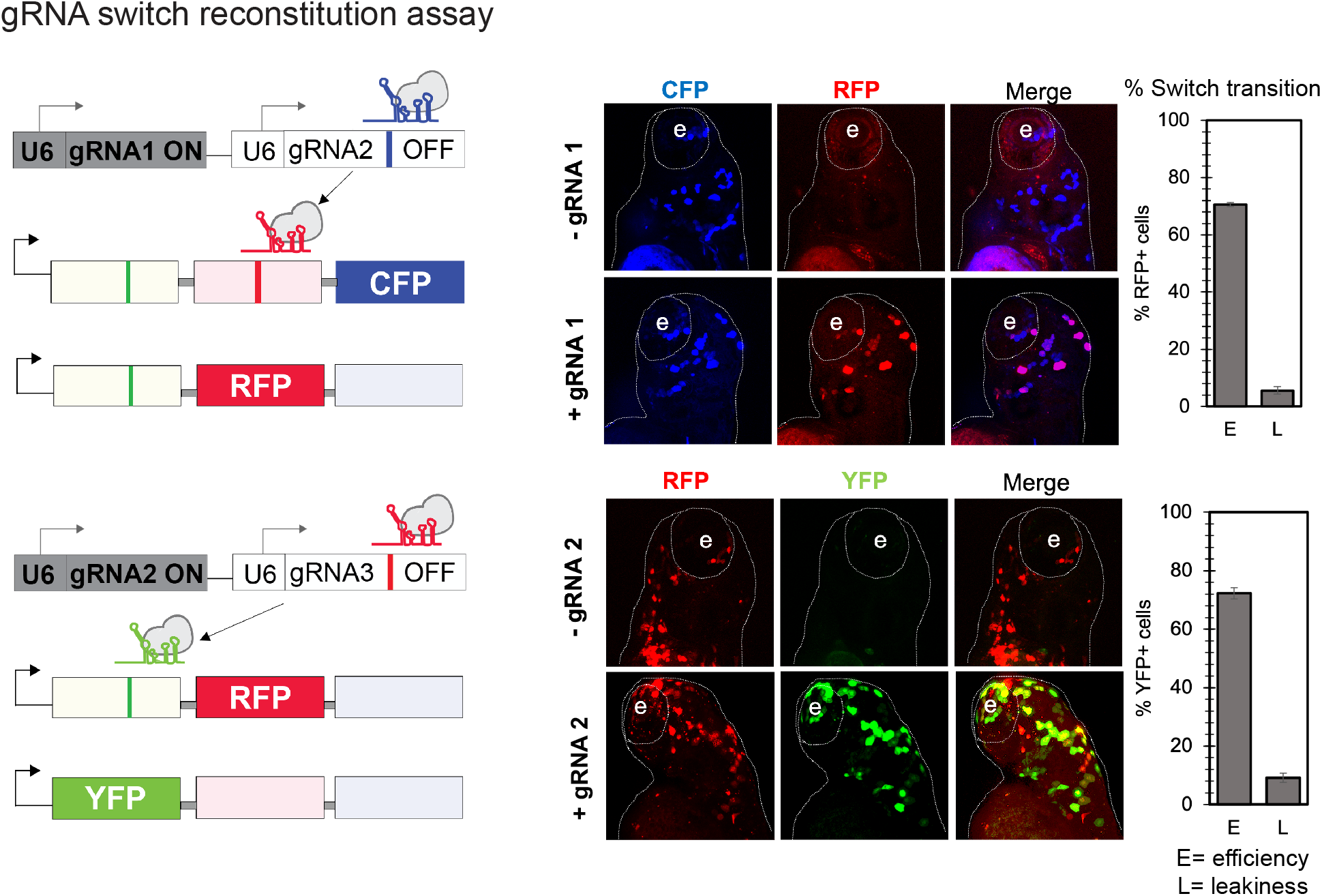
Different gRNA switches have similar reconstitution efficiency. Conditional gRNA switch transitions for two different target gRNA sequences, assessed by expression of fluorescent reporters in 3 day-post-fertilization (dpf) zebrafish larvae injected at 1-cell-stage. Plots indicate the mean ± SEM of the fluorescence for the reporter activated in the presence or absence of the corresponding gRNA, representing efficiency (E) or leakiness (L) of the transition, respectively (n=18 fish). Both gRNA switches show similar repair efficiency and leakiness, suggesting that keeping the conditional gRNA structure (repeat length, polyT and target orientation, see Fig. S1) is enough to maintain the gRNA switch properties even with different target sequences.

**Fig. S3.**
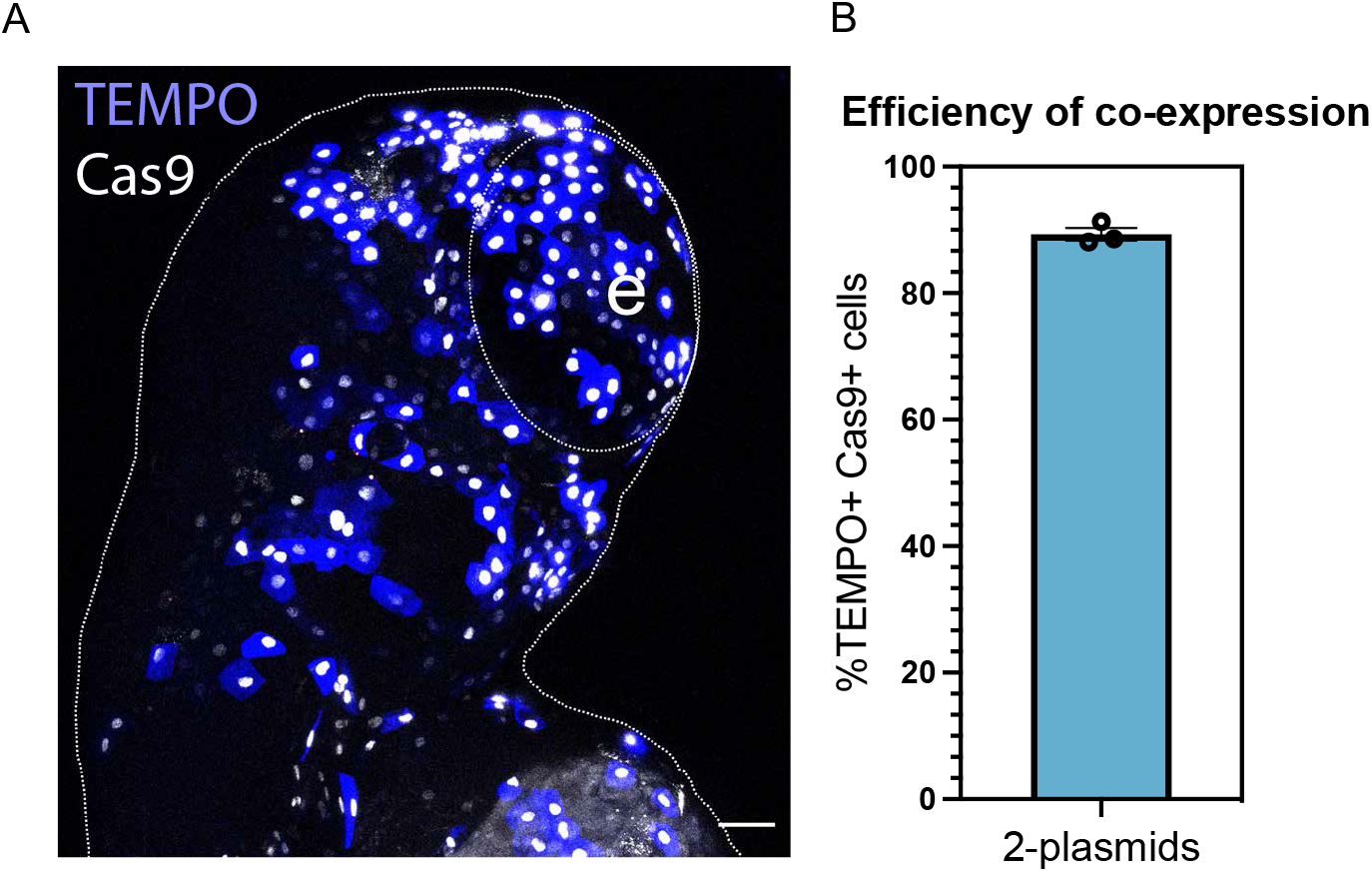
Co-expression of TEMPO and Cas9 plasmids in injected zebrafish is highly efficient. (**A**) Representative image of 3dpf zebrafish larvae co-injected with ubi:TEMPO (blue labelling) and ubi:Cas9 (white nuclear labelling) constructs (without gRNAs) at the 1-cell-stage. e =eye. (**B**) A high percentage of cells (>88%) co-express both constructs as seen in the plot (mean ± SEM, n=3 fish). Scale bar = 50 µm.

**Fig. S4.**
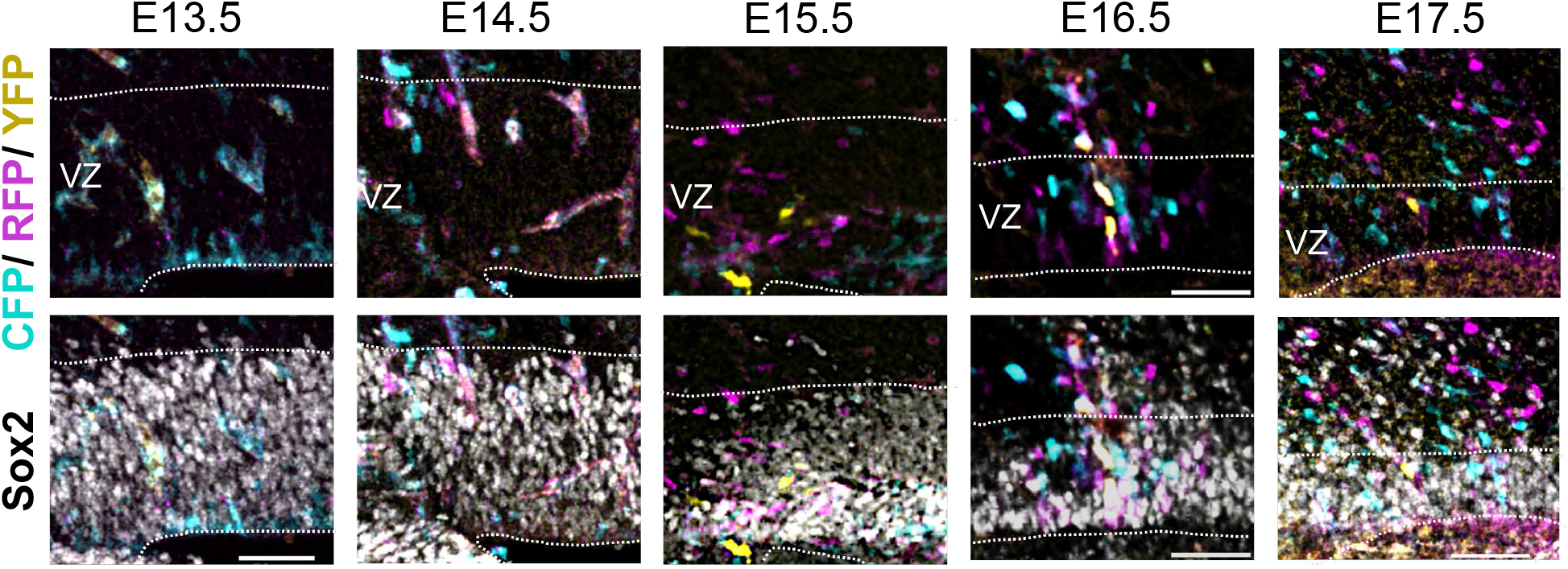
Sox2 co-labelling identifies TEMPO+ radial glial cells in the ventricular zone. Mouse brain cortices electroporated with TEMPO constructs at E12.5 and imaged at consecutive developmental stages. Co-labelling with Sox2 (lower panels) was used to identify TEMPO+ radial glial cells in the ventricular zone (VZ, dashed line contour). We distinguished Sox2 expression in the VZ from that of other adjacent cortical regions because of its higher levels of expression (*46*) and higher cell density in this region. (Sections E13.5-E15.5 and E17.5 are adjacent to the sections shown in Fig. 4B; section E16.5 is the same as shown in Fig. 4B, here showing Sox2 labelling as well). Scale bar = 50 µm.

**Fig. S5.**
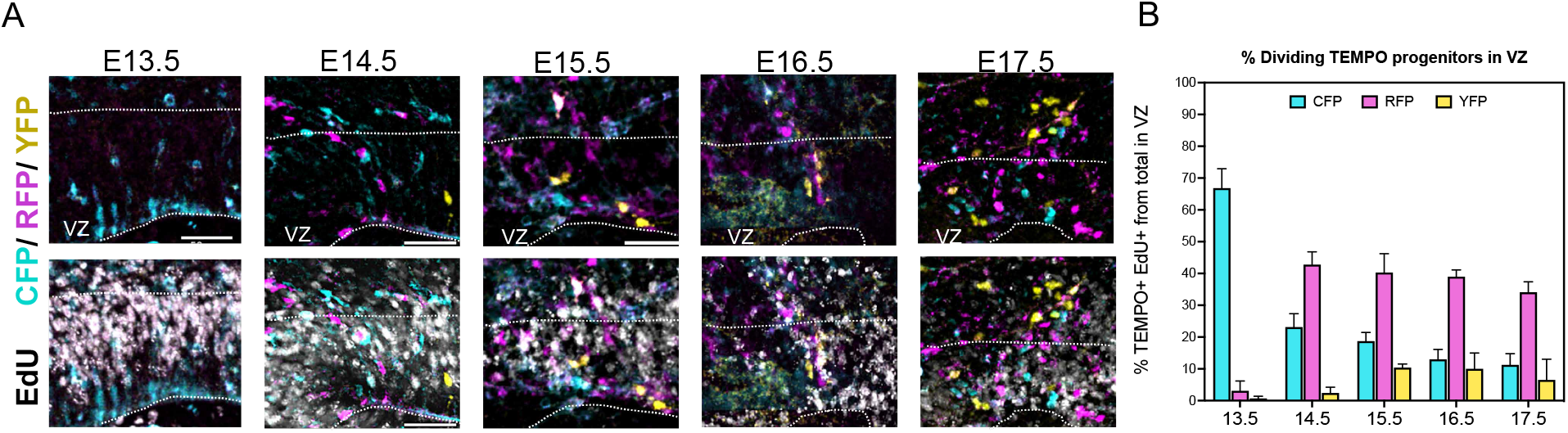
Dividing TEMPO+ progenitors decrease along time. **(A)** TEMPO+ cortices imaged at consecutive developmental stages after a 30-minute pulse of EdU to distinguish dividing progenitors. EdU co-labelling is shown in lower panels. (VZ= dashed area in A was determined by Sox2 co-labelling in adjacent sections, c.f. Fig. 4B and Fig. S4). Scale bar = 50 µm. (**B**) Color distribution of dividing TEMPO+ progenitors from total in VZ. The proportion of dividing progenitors, which can undergo reporter cascade transitions, decreases over time as seen in the plot (mean ± SEM, 3 independent experiments).

**Fig. S6.**
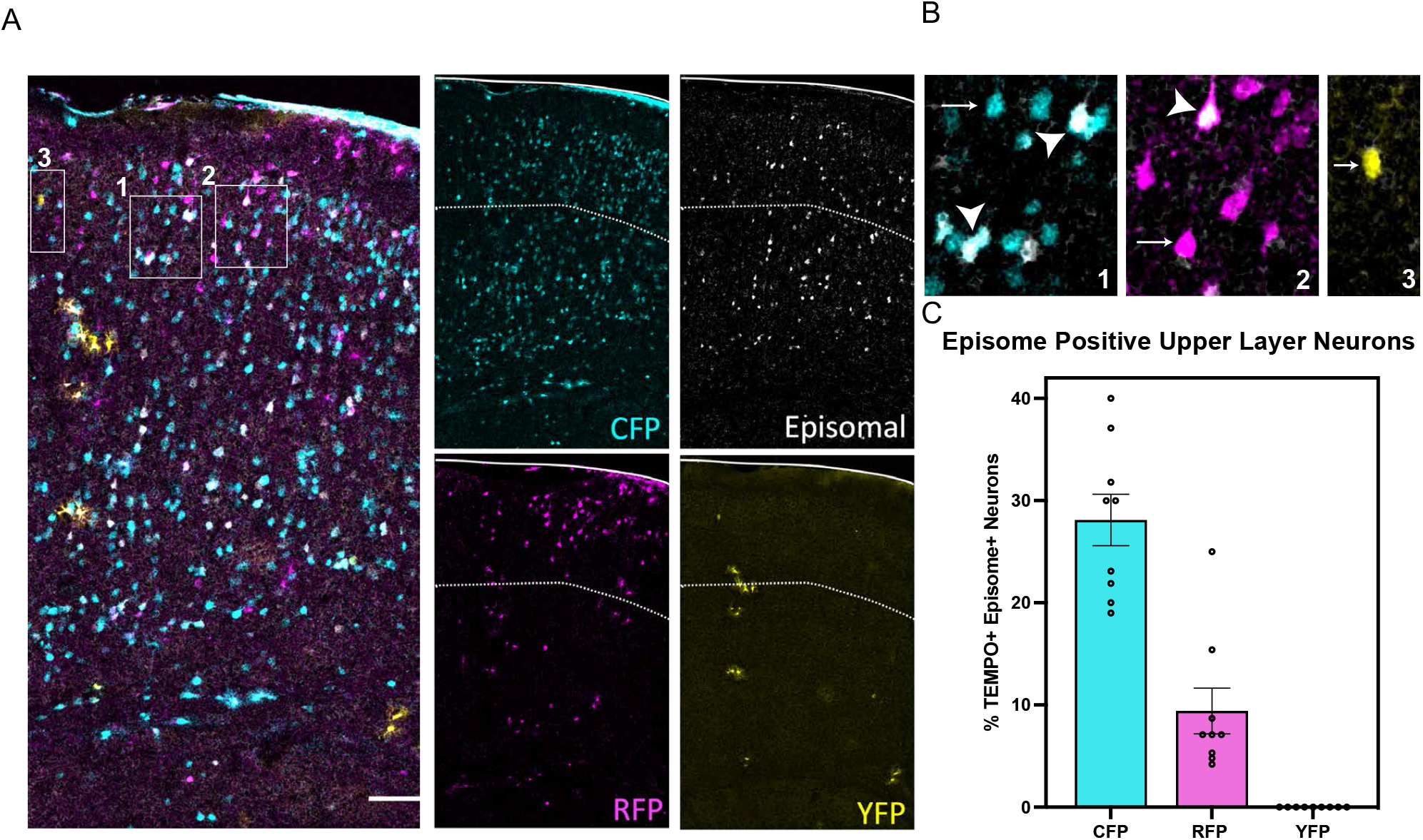
Low division capacity of a population of CFP+ progenitors contributing to upper layer neurons. **(A)** Postnatal P10 brain sections from mice electroporated with TEMPO integrative construct and Halotag episomal construct at E12.5. Scale bar = 100 µm. Insets in (**B**) show examples of neurons co-expressing TEMPO and Halotag which did not divide much after the electroporation, given they conserve the episomal plasmid (arrowheads in 1-2), and neurons only expressing TEMPO (arrows in 1-3), which might have lost the episomal plasmid after several rounds of cell division. (**C**) Higher proportion of CFP+ neurons in upper layers maintain episomal Halotag expression compared to RFP+ neurons, or YFP+ which do not express Halotag at all, revealing differences in their past proliferation history (mean ± SEM, n= 9 brain sections, 3 independent experiments).

**Fig. S7.**
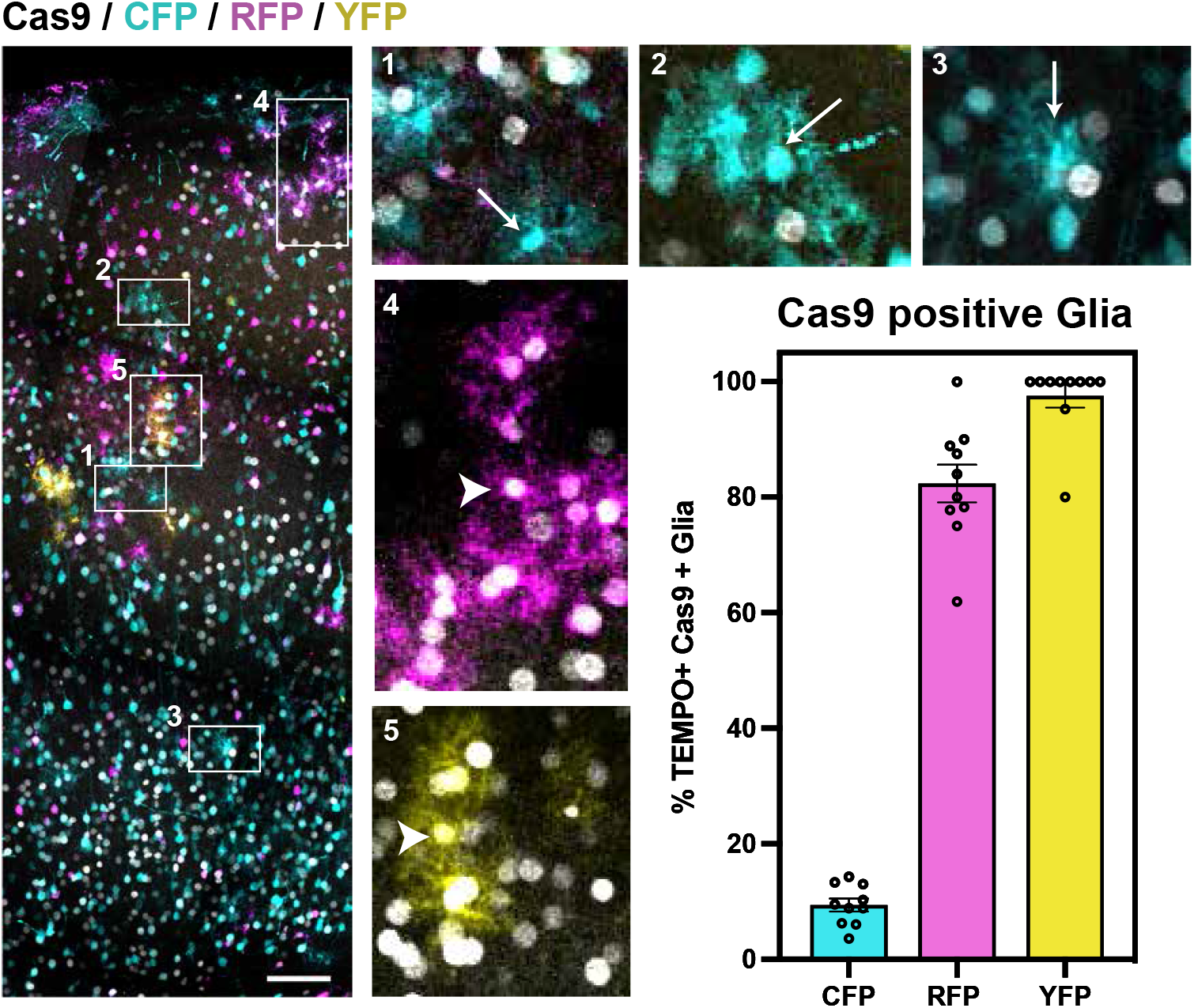
Analysis of the co-expression of Cas9 and TEMPO plasmids in glial cells. **(A)** Postnatal P10 brain sections from mice electroporated with TEMPO and Cas9-H2B-Halotag (gray nuclear labelling) constructs at E12.5. Scale bar = 100 µm. Insets reveal that most CFP+ astrocytes only express TEMPO but not Cas9-H2B-Halotag (arrows in 1-3), while most RFP+ and YFP+ astrocytes co-express both plasmids (arrowheads in 4 and 5), as seen in the plot (mean ± SEM, n= 10 brain sections, 3 independent experiments). These results show that most CFP+ glia observed did not transition in the reporter cascade because of the lack of Cas9.

**Fig. S8.**
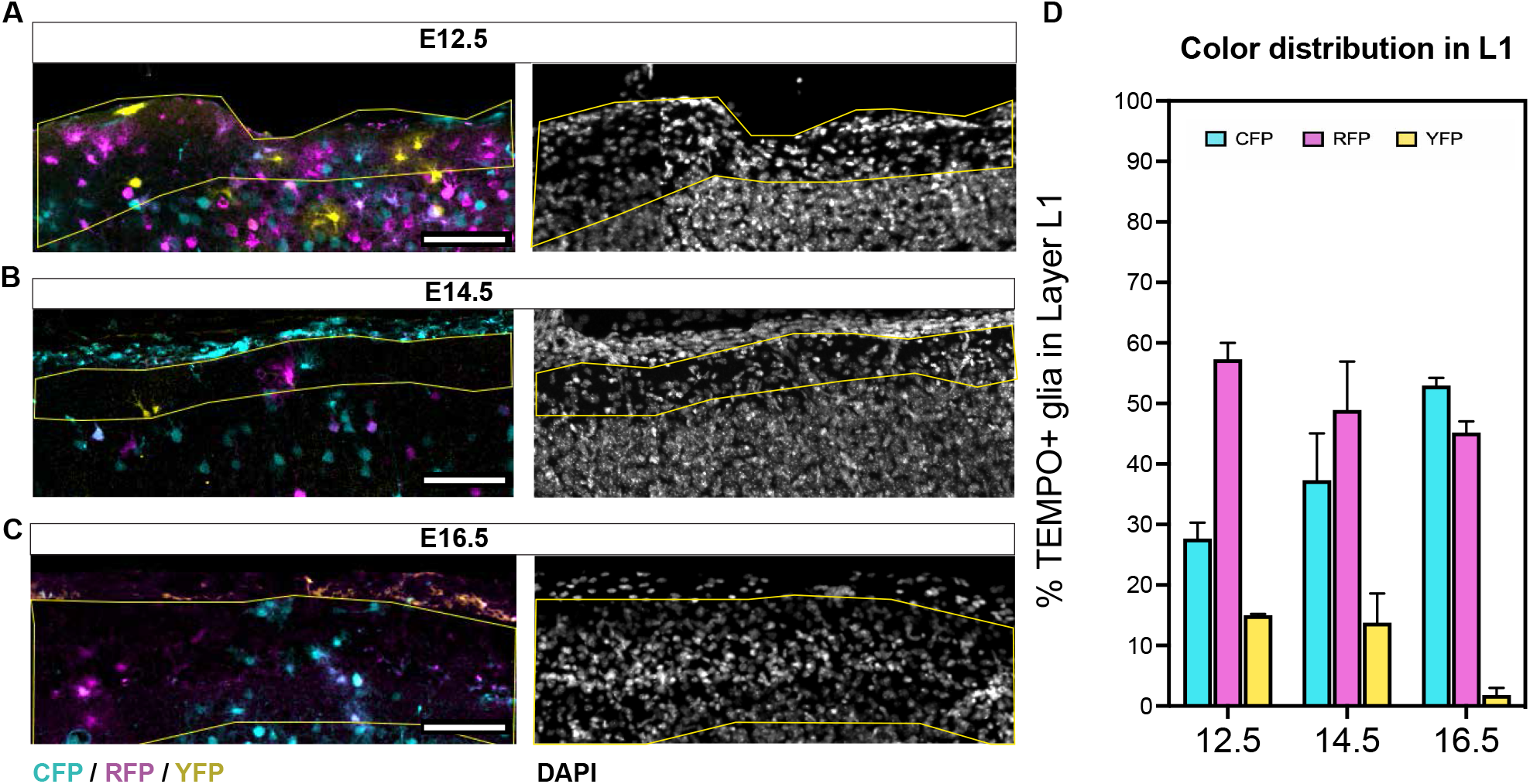
TEMPO color distribution in layer 1 (L1) astrocytes reveals differences in cascade progression along time. **(A-C)** Postnatal P10 brain sections from mice electroporated with TEMPO constructs at E12.5, E14.5 or E16.5 focusing on the most superficial cortical layer 1. DAPI labelling (right separated panels) was used to determine the area occupied by L1 (dashed yellow line). Scale bars = 100 µm. (**D**) TEMPO color progression in L1 astrocytes is lower in mice electroporated at later stages, especially after electroporation at E16.5. This suggests pial astrocytes do not divide much after E16.5 (mean ± SEM, n= 3 independent experiments).

**Fig. S9.**
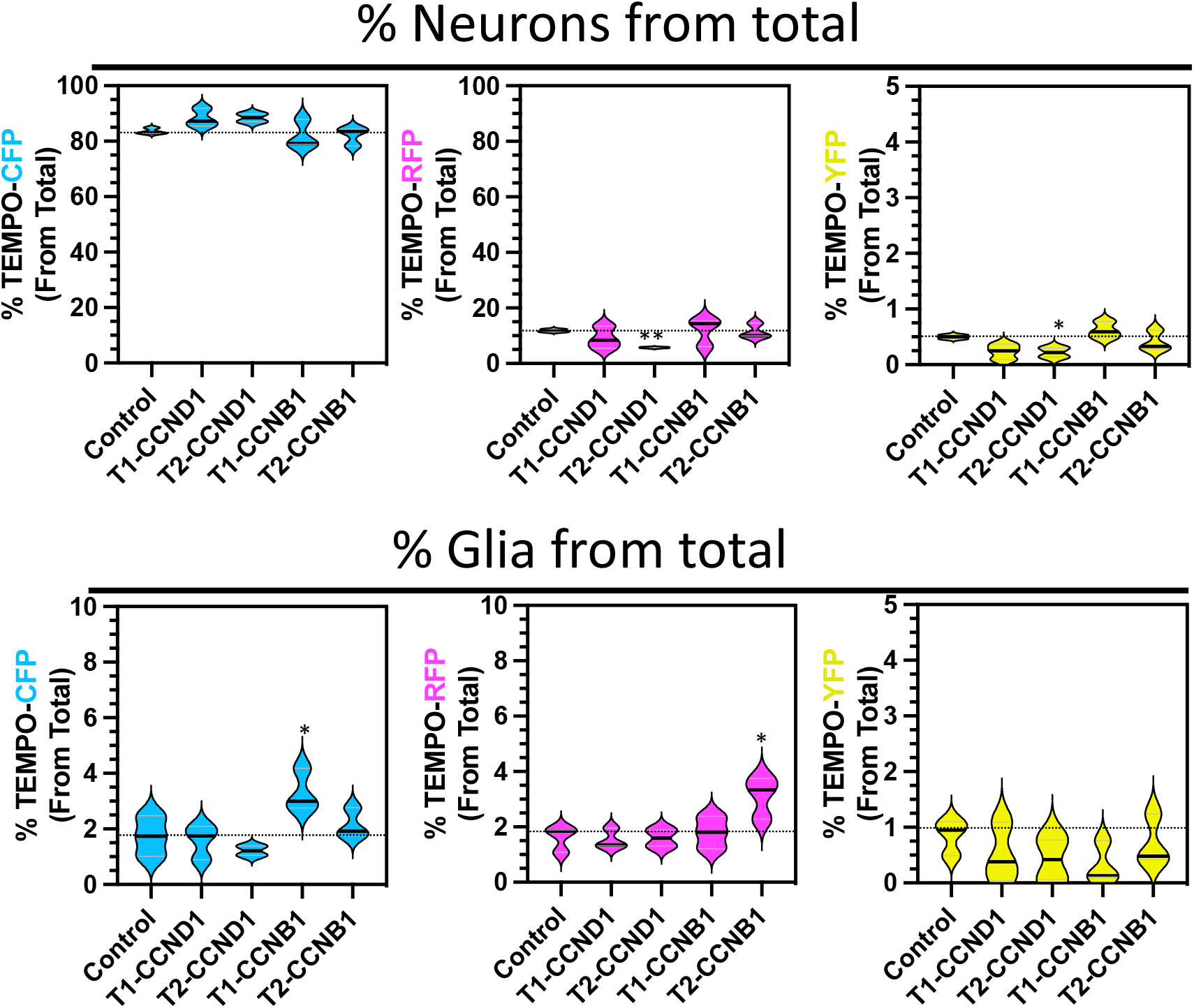
Proportions of cortical neurons and glia after temporal genetic manipulation of cell cycle regulators. **(A-C)** Postnatal P10 brain sections from mice electroporated with TEMPO-2.0 perturbation constructs at E12.5. Violin plots represent the total changes in neuron (upper panels) or glial cell numbers (lower panels) in control or manipulated samples. CCNB1, Cyclin B1. CCND1, Cyclin D1. We found a significant decrease in RFP+ (^**^p<0.01) and YFP+ (^*^p<0.05) neurons when overexpressing Cyclin D1 in the second (T2) temporal window (a two-tailed unpaired Student’s t test was used. n=3).

**Table S1.**
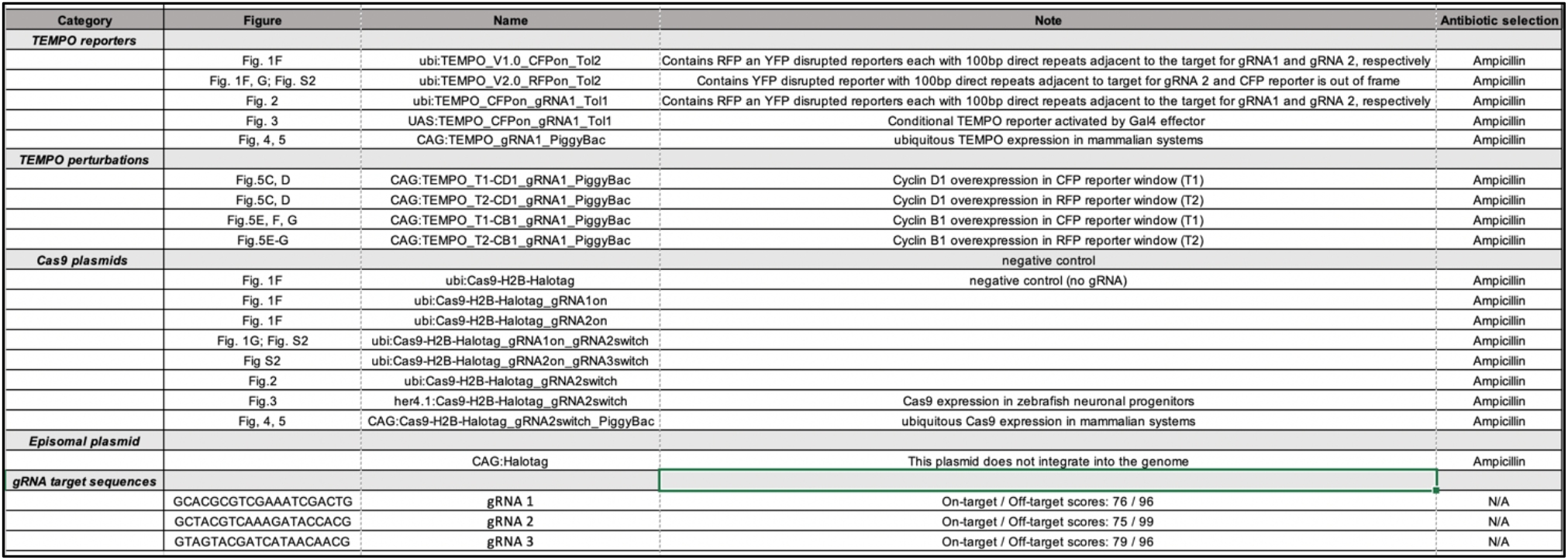
Plasmids used in the manuscript and gRNA sequences.

